# Decoding and modulating multiregional communication in the mood processing network

**DOI:** 10.1101/731547

**Authors:** Shaoyu Qiao, J. Isaac Sedillo, Kevin A. Brown, Breonna Ferrentino, Bijan Pesaran

## Abstract

Neural decoding and neuromodulation technologies hold great promise for treating mood and other brain disorders in next-generation therapies that manipulate functional brain networks. Here, we perform a novel causal network analysis to decode multiregional communication in the primate mood processing network and determine how neuromodulation, short-burst tetanic microstimulation (SB-TetMS), alters multiregional network communication. The causal network analysis revealed a mechanism of network excitability that regulates when a sender stimulation site communicates with receiver sites. Decoding network excitability from neural activity at modulator sites predicted sender-receiver communication while SB-TetMS neuromodulation specifically disrupted sender-receiver communication. These results reveal specific network mechanisms of multiregional communication and suggest a new generation of brain therapies that combine neural decoding to predict multiregional communication with neuromodulation to disrupt multiregional communication.

**One Sentence Summary:** Decoding and modulating multiregional network communication.

## Introduction

Brain-machine interfaces seek to achieve therapeutic effects by decoding recorded neural activity (*1, 2*). Neuromodulation technologies offer therapies by intervening to alter neural activity (*3*). The combination of neural decoding and neuromodulation technologies has broad and wide-ranging potential to treat refractory neurological and neuropsychiatric disorders (*1*). Both technologies can be performed using implantable devices to stimulate as in deep brain stimulation (*4*) and direct cortical surface stimulation (*5*) and record (*1*). Ultimately, next-generation neuromodulation-and-decoding-based therapies may be able to repair and even cure the disordered brain by recruiting neural plasticity.

Despite the therapeutic promise and recent advances, significant hurdles exist, in particular, for treating mood disorders. The primate mood network spans the orbitofrontal cortex (OFC), anterior cingulate cortex (ACC), dorsal prefrontal cortex (dPFC), striatum, amygdala (Amyg), hippocampus, nucleus accumbens (NAc), and ventral tegmental area. Functional interactions between these brain regions play a central role in regulating mood state variations (*6–9*) and modulating distributed computations in decision-making (*10, 11*). Dysfunction in the mood network alters motivation to guide behavior, underlies mood disorders such as major depression and anxiety (*12*), and often involves multiple brain regions distributed across the frontostriatal limbic network (*7, 12*). Neuromodulation has often been based on focal stimulation (*13–16*) and recent neural decoding approaches have decoded mood states to target neuromodulatory interventions (*6*). While clinical targets for therapeutic stimulation, such as for mood, are increasingly viewed within a circuit-based model (*17*) in which dysfunction arises from disordered multiregional communication, how specific mechanisms of multiregional communication can be decoded and manipulated is unknown.

Neuromodulation may suppress or facilitate activity at specific sites as well as weaken or strengthen multiregional communication. Disambiguating network mechanisms is a grand challenge in modern neuroscience. Communication is often estimated from correlations in activity at each site, or node (*18*), but correlations are confounded by changes in activity at each site and common inputs to both sites (*19*). To better identify network mechanisms, causal approaches based on simultaneous stimulation and recordings from multiple, interconnected regions are required but have been technically prohibitive in the awake primate brain. New approaches are therefore needed to identify, decode and manipulate multiregional communication to realize next-generation brain therapies.

Here, we address the need for novel neural decoding and neuromodulation-based therapies by analyzing multiregional communication. We present a neural decoding approach to test the network mechanism of action of a neuromodulatory intervention, short-burst tetanic microstimulation (SB-TetMS; ≤ 2 s, 100/200 Hz), by stimulating and recording across the large-scale mood processing network of two awake rhesus macaque monkeys. Importantly, our experiments analyze multiregional communication by extending the state-of-the-art with stimulation-recording coverage that spans OFC, dPFC, primary motor cortex (M1), parietal cortex, cingulate cortex, striatum, pallidum, and Amyg. We use the concept of neural excitability, which describes how each node responds to input according to the weight of an edge, to causally estimate multiregional communication. The resulting causal network analysis permits a rigorous dissection of network communication hypotheses. Our results provide a new basis for developing next-generation therapies in which neuromodulation specifically disrupts multiregional communication revealed by decoding network excitability.

### Targeting and sampling the mood processing network

**Figure 1A** illustrates the experimental design for decoding and altering multiregional communication. We first measure neural excitability by measuring neural responses across the network to a secondary intervention ― an isolated single microstimulation pulse. Decoding the neural responses is then used to identify modulators that predict network excitability moment-by-moment. We then deliver the primary intervention, a SB-TetMS pulse train, at sites in either gray matter or white matter and repeat the secondary intervention.

**Fig. 1.**
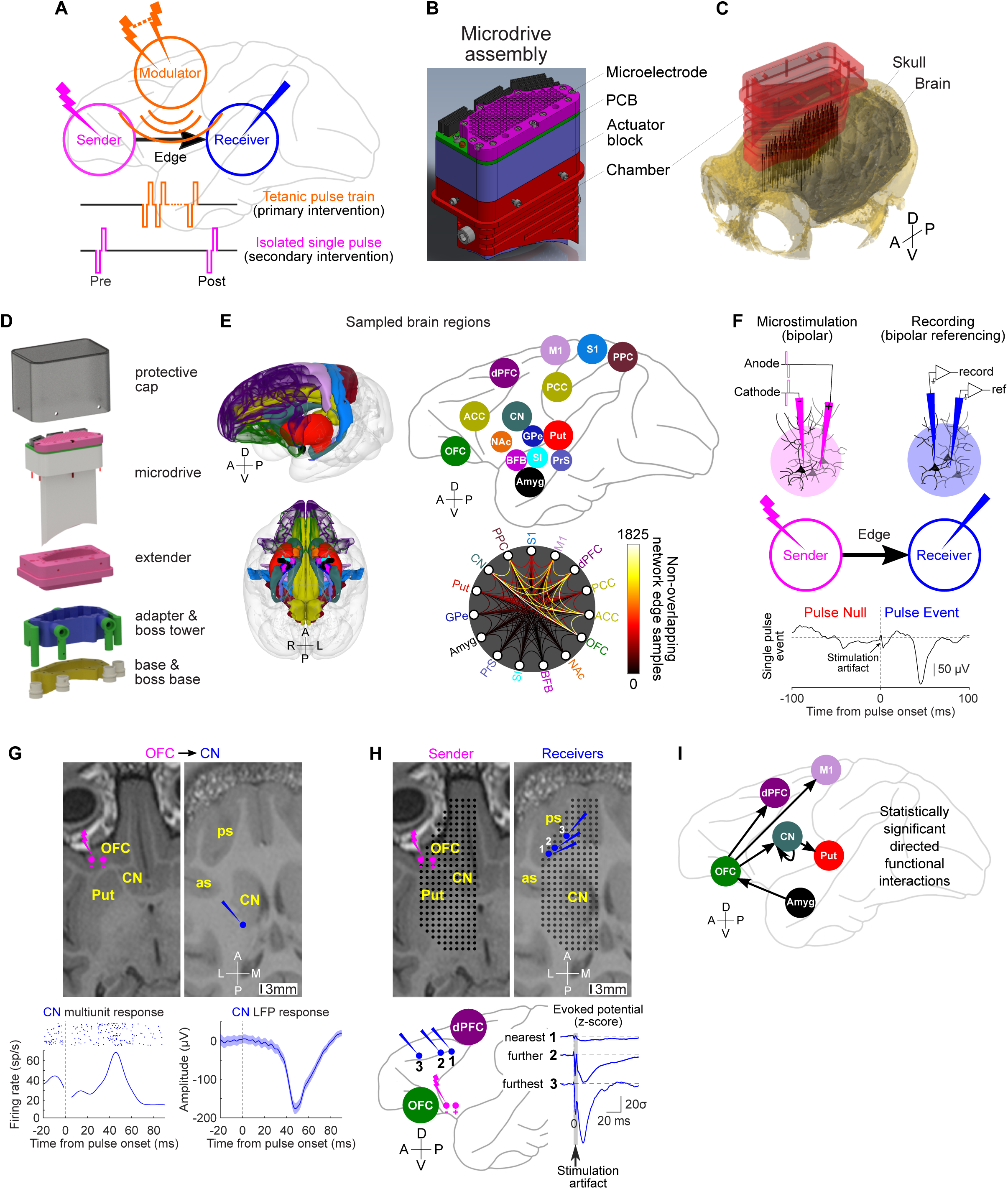
Chronic large-scale microdrive enables targeting and sampling the primate mood processing network. (**A**) Experimental design using multisite spatiotemporal patterned microstimulation to analyze neuromodulation hypotheses of network mechanisms of action of SB-TetMS-based intervention (primary) ― edge-modulation versus node-modulation. (**B**) Computer-aided design model of the customized large-scale microdrive assembly. (**C**) Co-registered chamber with MRI-based skull and brain surface reconstruction. Vertical black lines: Co-registered electrode tracks. (**D**) Computer-aided design model of modular chronic implant system. (**E**) Sampled brain regions (network nodes) illustrated using a rhesus macaque brain atlas (*37*) from the Scalable Brain Atlas. Non-overlapping network edge samples between sampled network nodes shown. (**F**) Single, bipolar, biphasic, charge-balanced microstimulation pulses at the sender to causally identify sender-receiver communication (black arrow). Example of CN receiver response to a single microstimulation pulse at OFC sender (30 μA, 100 μs/ph). (**G**) Single bipolar microstimulation pulses (10 μA, 100 μs/ph) at OFC sender drove both multiunit and LFP responses at CN receiver (response onset: ∼20 ms, response peak: ∼46 ms after pulse onset). Data around the pulse onset omitted for multiunit activity analysis due to stimulation artifact. Shading: +/-1 s.e.m. (*n* = 52). (**H**) Single bipolar microstimulation pulses (50 μA, 100 μs/ph) at OFC sender revealed significant focal evoked LFP responses in principal sulcus (ps) distant from the stimulating site, not nearby. Horizontal MRI slice at OFC stimulating sites (Left; magenta dots) and ps recording sites (Right; blue dots). (**I**) Network diagram showing statistically significant sender-receiver communication. Anterior (A), posterior (P), left (L), right (R), medial (M), lateral (L), dorsal (D), and ventral (V) directions shown.

To decode multiregional communication and analyze neuromodulation across the mood processing network, we developed a novel implantable microdrive to target and sample a large-scale mood processing network for causal network analysis (**Fig. 1, B-D, and fig. S1**). With this device, we successfully targeted and repeatedly sampled a network spanning 15 cortical and subcortical brain regions. The sampled network included seven cortical regions: OFC, ACC, posterior cingulate cortex (PCC), dPFC, M1, primary sensory cortex (S1), and posterior parietal cortex (PPC); and eight subcortical regions: caudate nucleus (CN), putamen (Put), globus pallidus external (GPe), Amyg, presubiculum (PrS), substantia innominata (SI), basal forebrain nucleus (BFB), and NAc.

The causal network analysis let us study neural interactions between sets of recording and stimulation sites located at nodes in the network. Across all 15 nodes, we sampled from a grand total of 4658 sites in Monkey M and 4897 sites in Monkey A (**Fig. 1E**). Across both monkeys, we obtained 15,734 non-overlapping samples from 21 cortico-cortical edges, 11,276 non-overlapping samples from 56 cortical-subcortical edges, and 1087 non-overlapping samples from 28 subcortical-subcortical edges (**Fig. 1E**). We targeted our analysis to 9548 samples from the mood processing network connecting OFC, ACC, dPFC, CN, Put, and Amyg.

### Causal network analysis reveals multiregional communication

We causally estimated multiregional communication with a secondary intervention delivering a single, low-amplitude, bipolar, biphasic, charge-balanced microstimulation pulse at a given node (sender) while recording evoked responses at other sampled nodes (receiver). Responses to isolated bipolar microstimulation pulses appear to reflect responses by neurons in the receiver due to synaptic inputs driven by stimulation at the sender (sender→receiver; **Fig. 1F**). Consistent with this interpretation, single bipolar microstimulation pulses drove spiking activity from neurons in the receiver, as well as the evoked LFP activity (**Fig. 1G**). Thus, bipolar re-referenced LFP responses reflected a local neuronal source. Moreover, analyzing the spread of signals from the sender site to the receiver sites confirmed that using bipolar microstimulation and recording configurations revealed focal responses at receiver sites distant to the sender (**Fig. 1H**). Comparison with control recordings during monopolar microstimulation (**fig. S2**) confirmed that bipolar microstimulation was less likely to contaminate receiver site responses due to current spread.

To detect pulse-evoked responses, we used an optimal signal detection algorithm (*20*), which we term stimAccLLR. Across 12,721 pairs of sites tested in 132 network edges we causally sampled, we identified and characterized 78 significant directed functional interactions (sender-receiver communication) in 21 network edges (**figs. S3 and S4; tables S1 and S2**). On average, we identified sender-receiver communication between a pair of nodes with probability of 0.61%. Since not all pairs of nodes are expected to functionally interact, we tested whether some pairs of nodes interacted more than others (**table S2**). A subset of edges revealed significant directed functional interactions (*P* < 0.05, binomial test; **Fig. 1I**): OFC→CN (1.16%: 14/1212 samples, *P* = 0.009), OFC→dPFC (1.16%: 9/779 samples, *P* = 0.027), OFC→M1 (6%: 6/100 samples, *P* = 3.2×10^−5^), CN→Put (4.46%: 5/112 samples, *P* = 5.4×10^−4^), and Amyg→OFC (10.17%: 6/59 samples, *P* = 1.5×10^−6^). We also observed significant interactions within a single node, such as CN→CN (1.23%: 7/568 samples, *P* = 0.035). Thus, the causal network analysis defined a cortical-subcortical limbic network (OFC-dPFC-M1-CN-Put-Amyg). We then set out to determine how SB-TetMS neuromodulation alters or modulates multiregional communication and how such communication can be decoded from activity at modulator sites.

### SB-TetMS suppresses excitability across the network

We first asked whether the network mechanism of SB-TetMS neuromodulation is consistent with either or both of the edge- and node-modulation hypotheses. We tested the edge-modulation hypothesis by asking whether stimulating the modulator site alters the strength of a sender-receiver communication (**Fig. 2A**). We did so by first probing the sender-receiver pair before delivering SB-TetMS at a modulator site that, critically, was not at the sender or the receiver site, and then probing the sender-receiver pair again immediately after SB-TetMS (median-T_post_ = 50 ms). **Figure 2B** illustrates potential outcomes for the edge-modulation hypothesis as a suppression or facilitation of neural excitability as revealed by the pulse-triggered evoked response in Pre-tetanic and Post-tetanic epochs. We also tested the node-modulation hypothesis for neural excitability by asking whether TetMS-modulator alters activity within the receiver node alone. **Figure 2C** illustrates potential outcomes for the receiver node-modulation hypothesis.

**Fig. 2.**
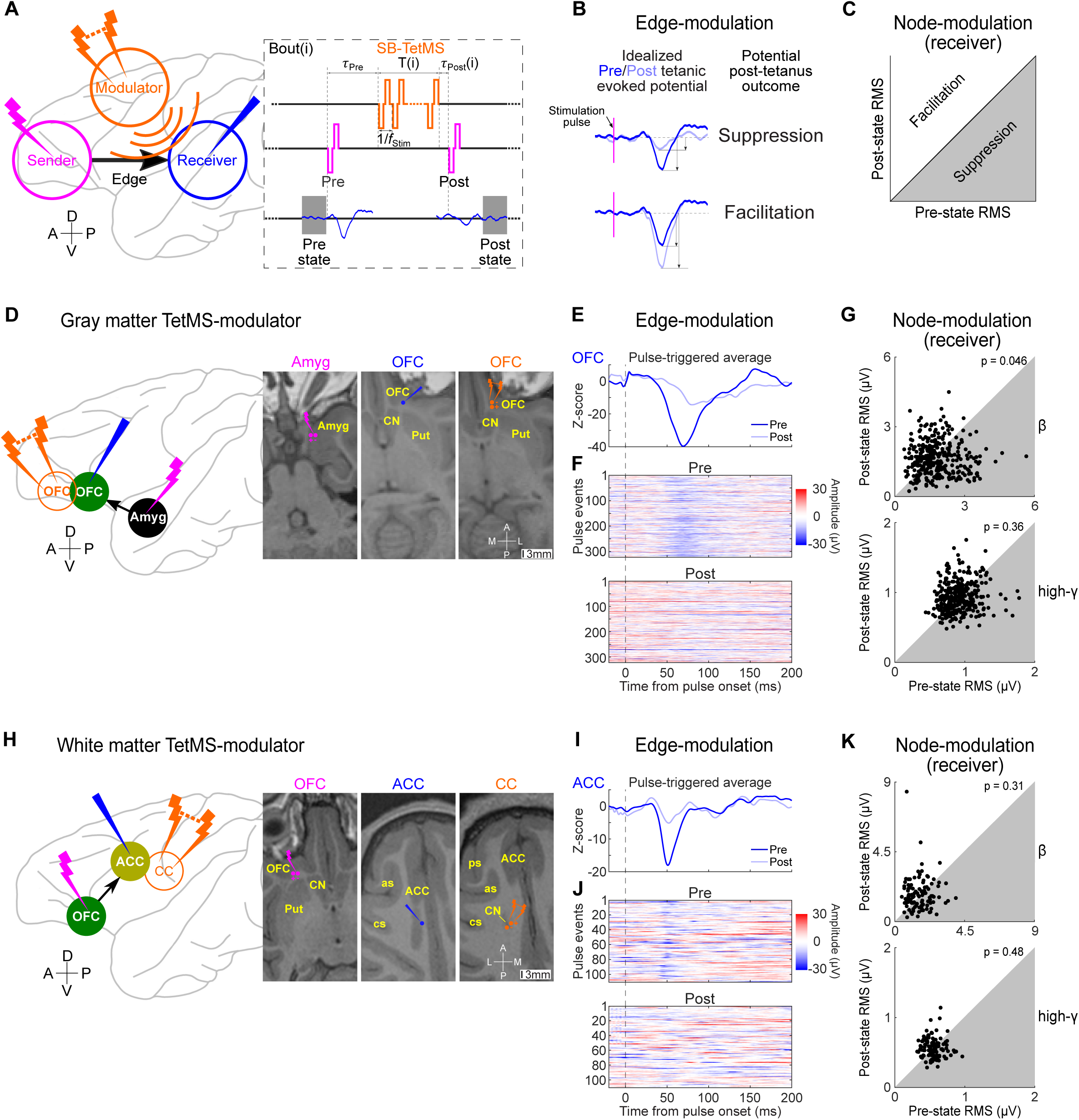
SB-TetMS suppresses excitability of sender-receiver communication. (**A**) Illustration of how TetMS-modulator alters the edge weight and the receiver node activity. For each stimulation bout, T_Pre_ is the post Pre-tetanic pulse epoch before the TetMS onset, T is the duration of TetMS with a given frequency (f_stim_), and T_Post_ is the latency between TetMS offset and Post-tetanic single pulse onset. The gray shaded rectangles are the 100-ms epochs for computing Pre-state RMS and Post-state RMS of the neural activity recorded at the receiver, respectively. RMS denotes the root mean square. (**B**) Illustration of testing edge-modulation hypothesis by comparing Pre-tetanic and Post-tetanic evoked potentials. (**C**) Illustration of testing receiver node-modulation hypothesis by comparing the RMS power of Pre-state and Post-state neural activity at the receiver. (**D**) OFC TetMS-modulator (T = 50-500 ms, f_stim_ = 100 Hz, 30 μA, 100 μs/ph; orange dots) suppressed Amyg→OFC excitability revealed by evoked OFC receiver (blue dot) response to single pulses (40 μA, 100 μs/ph; *T*_Pre_ = 500 ms and *T*_Post_ = 25-100 ms) delivered at an Amyg sender (magenta dots). (**E**) Z-scored pulse-triggered average OFC receiver responses (*n* = 322) to Pre-tetanic and Post-tetanic single pulses respectively (Amyg→OFC). (**F**) Evoked OFC receiver responses pulse-by-pulse during Pre-tetanic and Post-tetanic epochs, respectively. (**G**) Scatter plots of Pre-state RMS versus Post-state RMS of OFC receiver β and high-γ neural activity, respectively. (**H**) Corpus callosum (CC) TetMS-modulator (T = 250 ms, f_stim_ = 200 Hz, 20 μA, 100 μs/ph; orange dots) suppressed OFC→ACC excitability revealed by evoked ACC receiver (blue dot) response to single pulses (30 μA, 100 μs/ph; *T*_Pre_ = 500 ms and *T*_Post_ = 0, 50 ms) delivered at an OFC sender (magenta dots). as: arcuate sulcus; cs: central sulcus; ps: principal sulcus. (**I**) Z-scored pulse-triggered average ACC receiver responses (*n* = 110) to Pre-tetanic and Post-tetanic single pulses, respectively (OFC→ACC). (**J**) Evoked ACC receiver responses pulse-by-pulse during Pre- and Post-tetanic epochs, respectively. (**K**) Scatter plots of Pre-state RMS versus Post-state RMS of ACC receiver β and high-γ neural activity, respectively. Anterior (A), posterior (P), lateral (L), medial (M), dorsal (D), and ventral (V) directions shown.

**Figure 2D–2G** presents a gray matter TetMS-modulator. Here, we stimulated an Amyg sender and observed OFC receiver responses (**Fig. 2E**). The evoked response was visible pulse-by-pulse (**Fig. 2F and fig. S5, A-C**). SB-TetMS delivered at an OFC modulator site (**Fig. 2D**) suppressed Post-tetanic OFC receiver responses (**Fig. 2E**). The OFC TetMS-modulator also slightly decreased the OFC receiver β power (*P* = 0.046, Wilcoxon rank sum test; **Fig. 2G**). In comparison, SB-TetMS did not significantly change OFC receiver high-γ power (*P* = 0.36; **Fig. 2G**). Results for this Amyg-sender, OFC-receiver, and OFC-TetMS-modulator are consistent with both edge- and node-modulation mechanisms of neuromodulation.

**Figure 2H–2K** presents a white matter TetMS-modulator. Here, we stimulated an OFC sender and observed ACC receiver responses pulse-by-pulse (**Fig. 2, I and J and fig. S5, D-F**). Following SB-TetMS delivered to the corpus callosum (CC) (**Fig. 2H**), ACC receiver responses were suppressed (**Fig. 2I**). The CC TetMS-modulator did not significantly change the ACC receiver β (*P* = 0.31) nor high-γ power (*P* = 0.48) (**Fig. 2K**). Hence, this white matter TetMS-modulator reveals edge-modulation but not node-modulation.

We performed SB-TetMS across eight gray and white matter sites (**fig. S3 and table S1**) and tested 62 edge samples. Gray and white matter SB-TetMS showed clear edge modulation and suppressed excitability across all identified network edge samples except two that we could not measure due to stimulation artifacts (60/62 samples, **table S1**). Receiver node-modulation was more mixed. Gray matter TetMS-modulators significantly suppressed receiver node β power (11/40 nodes, *P* = 2.55×10^−6^, binomial test) but did not significantly change high-γ power (3/40 nodes, *P* = 0.19), while the white matter TetMS-modulator did not significantly suppress either (β: 1/20 nodes, *P* = 0.38; high-γ: 3/20 nodes, *P* = 0.06).

Therefore, SB-TetMS exerts a widespread neuromodulatory influence and overwhelmingly acts to suppress neural excitability across the mood processing network, temporarily shutting down multiregional communication. Gray matter SB-TetMS is more specific and also modulates β-power at the receiver node.

### Decoding sender-receiver communication from network excitability

Since SB-TetMS overwhelmingly modulates neural excitability across the network, excitability may reflect a network mechanism and involve other brain sites whose activity modulates sender-receiver communication (**Fig. 3A**). Therefore, activity at other network sites, i.e. modulators, may be associated with strengthening or weakening the receiver response to sender synaptic input pulse-by-pulse. If so, decoding sender-receiver communication may be possible by decoding modulator neural activity to predict receiver responses and, by doing so, reveal how the modulator alters sender-receiver excitability-based communication.

**Fig. 3.**
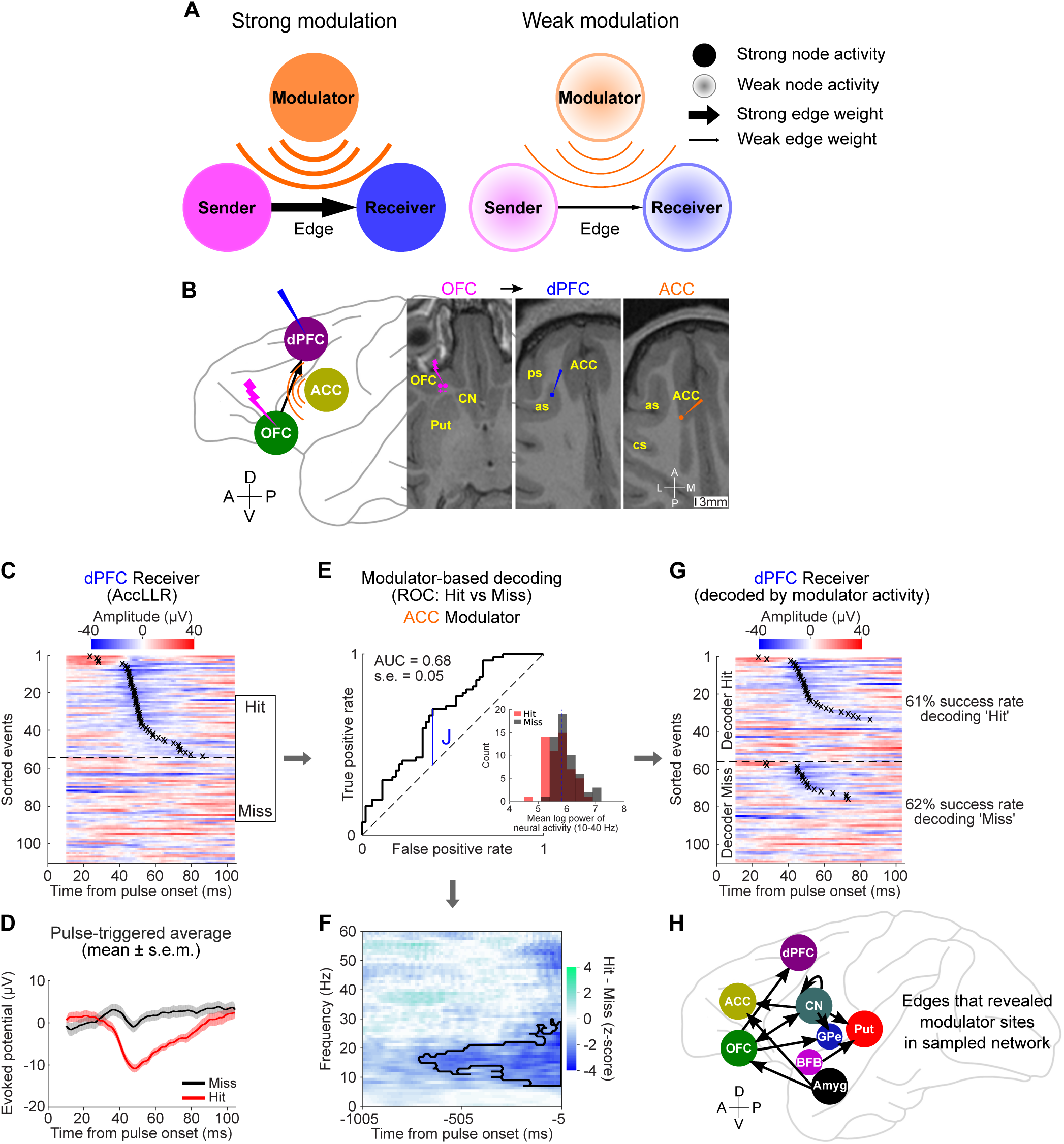
Modulator activity covaries with network excitability. (**A**) Schematic of strong and weak modulation of sender-receiver network excitability by modulator sites. (**B**) OFC→dPFC communication modulated by an ACC modulator (as: arcuate sulcus; ips: intraparietal sulcus). (**C**) Evoked dPFC receiver responses sorted by pulse-response latency (crosses). 49% “Hit” events (54/110), mean latency = 43 ms. (**D**) Pulse-triggered average evoked responses for ‘Hit’ (red) and ‘Miss’ events (black). Shaded: +/-1 s.e.m. (**E**) ROC analysis for decoding dPFC receiver events from ACC modulator (10-40 Hz power; FDR-corrected *P* = 0.0367). J: Youden’s index. Dashed line: chance level. Solid blue line: threshold for decoding, the dashed blue line on the histogram of ACC modulator power for ‘Hit’ (red) and ‘Miss’ (black) dPFC receiver events. (**F**) Modulation spectrograms of ACC modulator activity before Pre-tetanic pulse, computed as z-score magnitude of power difference between ‘Hit’ and ‘Miss’ events in dPFC receiver responses. Significance contour from permutation test (cluster-corrected; *n* = 10,000, *P* < 0.05, two-tailed test). (**G**) Decoding performance. Waveforms classified into ‘Decoder Hit’ and ‘Decoder Miss’ categories at the dPFC receiver from the ACC modulator activity. In each category, waveforms sorted by pulse-response latency (crosses). (**H**) Network diagram of modulated sender-receivers. Anterior (A), posterior (P), lateral (L), medial (M), dorsal (D), and ventral (V) directions shown.

**Figure 3B–4D** presents example OFC→dPFC communication. We detected the response to each Pre-tetanic pulse (**fig. S6**). Impressively, along with ‘Hit’ events, we observed ‘Miss’ events when the receiver did not respond to sender stimulation (**Fig. 3, C and D**). Of all edge samples identified, the missing responses matched the failure of sender-receiver communication following SB-TetMS at TetMS-modulator sites. Multiunit activity, when observable, was also strongly driven in ‘Hit’ events, but not in ‘Miss’ events (**fig. S7**). Unsupervised receiver response clustering provided convergent evidence for ‘Hit’ and ‘Miss’ events (**fig. S8**), consistent with the action of a modulatory network mechanism.

**Fig. 4.**
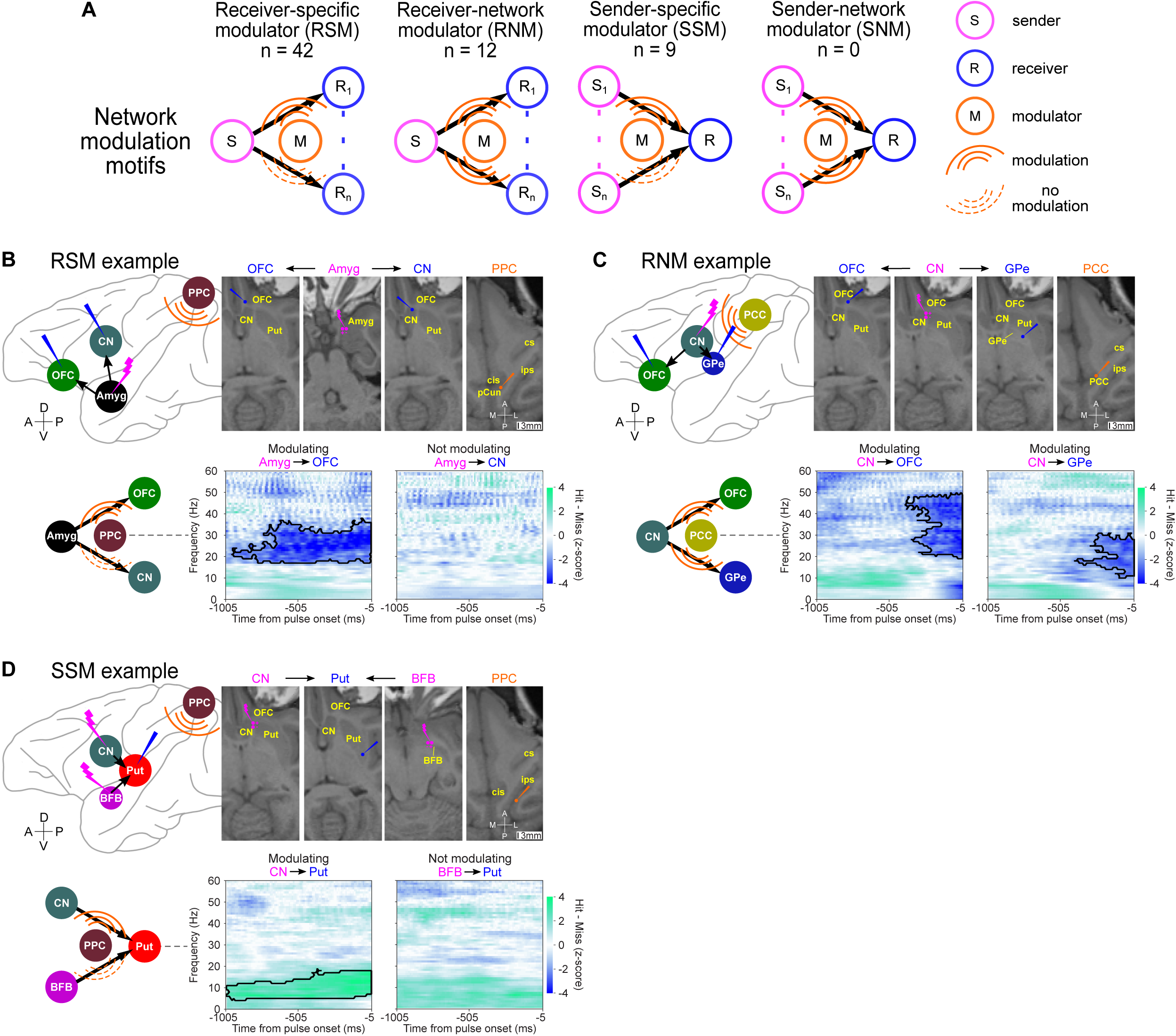
Network modulation motifs. (**A**) RSM, RNM, SSM, and SNM motifs. (**B**) RSM example: PPC modulator site in precuneus (pCun) predicted Amyg→OFC excitability (AUC = 0.63, s.e. = 0.03, FDR-corrected *P* = 0.019) not Amyg→CN excitability (AUC = 0.52, s.e. = 0.033, FDR-corrected *P* = 0.57). cs: central sulcus; ips: intraparietal sulcus; cis: cingulate sulcus. (**C**) RNM example: posterior cingulate cortex modulator predicted CN→OFC excitability (AUC = 0.63, s.e. = 0.032, FDR-corrected *P* = 0.0014) and CN→GPe excitability (AUC = 0.61, s.e. = 0.032, FDR-corrected *P* = 0.024). (**D**) SSM example: PPC modulator site in supramarginal gyrus predicted CN→Put excitability (AUC = 0.6, s.e. = 0.039, FDR-corrected *P* = 0.049) not BFB→Put excitability (AUC = 0.51, s.e. = 0.032, FDR-corrected *P* = 0.49). Modulation spectrogram contours show significance (permutation test, cluster-corrected; *n* = 10,000, *P* < 0.05, two-tailed test). Anterior (A), posterior (P), lateral (L), medial (M), dorsal (D), and ventral (V) directions shown.

To identify the modulatory network mechanism, we then built a neural decoder to decode sender-receiver communication. We specifically decoded modulator neural activity to predict ‘Hit’ and ‘Miss’ events using simultaneously recorded neural activity before each Pre-tetanic pulse. We focused on decoding activity between 10-40 Hz (**Fig. 3E**) as it reflects large-scale brain network mechanisms (*21*). **Figure 3F** shows increased ACC modulator power (10-30 Hz) significantly predicted OFC→dPFC communication (Hits: 61% Misses: 62%; **Fig. 3G**). From 62 identified sender-receiver pairs, we identified 63 modulator sites (1797 sites tested, *P* = 0.0022, binomial test) from 20 samples across 11 network edges (**Fig. 3H and tables S3–S5**). Thus, neural excitability can be decoded from neural activity at modulator sites, consistent with network excitability mechanism for sender-receiver communication.

### Sender-receiver-modulator network motifs

The neural decoding results suggest that multiregional communication across the network is dynamically orchestrated by modulator activity. If so, modulators should reveal specific motifs of network modulation with different mechanistic implications (**Fig. 4A**). If the mechanism involves receiver-input-specific modulation, when a sender projects to multiple receivers, modulator activity should only covary with excitability for a specific sender-receiver edge (receiver-specific modulator, RSM, motif). In contrast, if the modulation mechanism is shared across multiple projections, modulator activity should covary with excitability across multiple sender-receiver edges (receiver-network modulator, RNM, motif). Similarly, when multiple senders project to a single receiver, a sender-specific modulator (SSM) should only covary with excitability across a specific sender-receiver edge (SSM motif) whereas mechanisms of modulation shared across multiple projections should be revealed by a sender-network modulator (SNM) motif.

**Figure 4B** presents an RSM motif example. Increased power of a PPC modulator baseline activity in 20-40 Hz significantly predicted missing responses at an OFC receiver from an Amyg sender (Amyg→OFC), whereas the PPC modulator baseline activity did not modulate excitability of a CN receiver from the same Amyg sender (Amyg→CN). **Figure 4C** presents an RNM motif example. Increased power of a PCC modulator baseline activity in 20-50 Hz significantly predicted missing responses at an OFC receiver from a CN sender (CN→OFC), while increased power of the same PCC modulator baseline activity in 10-30 Hz also significantly predicted missing responses at the GPe receiver from the same CN sender (CN→GPe). **Figure 4D** presents an SSM motif example. Increased power of a PPC modulator baseline activity in 5-20 Hz slightly predicted hitting responses at a Put receiver from a CN sender (CN→Put), whereas the same PPC modulator baseline activity did not modulate excitability of the Put receiver from a BFB sender (BFB→Put). Thus, sender-receiver excitability-based communication can be modulated in an input-specific manner as well as by mechanisms shared across distributed networks.

Of 63 modulator sites identified, we observed input-specific mechanisms more often than shared mechanisms of modulation (42 RSM, 12 RNM, 9 SSM and no SMM motifs; **tables S3–S5**). Overall, we found PFC→basal ganglia (BG) and PFC→PFC communication were mainly modulated by other sites in cortex, while BG/Amyg→PFC and BG→BG communication were modulated by more widespread sites across cortex and BG.

## Discussion

Here, we present a large-scale causal network analysis to decode multiregional communication based on network excitability, and identify how SB-TetMS alters communication across the projections connecting OFC, ACC, dPFC, the motor cortices, CN, Put, and Amyg within the mood processing network. Our work shows how to go beyond focal electrical stimulation (*13–16*) by considering brain network communication. We reveal sender-receiver modulation and network excitability-based communication to show that network modulators regulate multiregional communication moment-by-moment. Our finding that neuromodulation perturbs network mechanisms regulating information flow between brain regions has major implications for understanding and treating functional brain networks. By showing that neural decoders can track variations in network excitability that underlie multiregional communication, and by showing that neuromodulation specifically alters sender-receiver communication, we demonstrate how next-generation brain therapies featuring excitability-based stimulation-recording neural interfaces can be developed to treat disordered brain networks.

Mood processing depends on a complex, distributed network whose interconnections maintain functional organization across multiple cortical-BG pathways (*22*). Existing approaches to reveal functional network dynamics in the primate mood and reward circuit are mainly based on functional MRI (fMRI) data (*8, 23*) which lacks spatiotemporal resolution and is relatively indirect. The large-scale microdrive approach used here provides more direct electrophysiological evidence to reveal how brain dynamics feature a rapid and precise mechanism of excitability-based modulation that allows selective communication between networks of senders and receivers. Synthesizing evidence from the sender-receiver-modulator motifs we observed across the cortical-BG pathway reveals that modulation of the BG responses to cortical inputs is primarily cortical in origin, specifically modulated by OFC, ACC, and dPFC. The BG output, measured by cortical responses to BG inputs, on the other hand, reflects modulation by many regions in both cortex and BG. This finding is likely due to how indirect pathways involving cortico-thalamic and BG-thalamic projections link BG to cortex (*22*). In both cases, modulator activity especially within the β-frequency band reveals a form of network gating that modulates sender-receiver communication, suppressing interactions momentarily in time. The network modulation motifs we observed also provide a new framework for studying value computation in the reward and decision-making circuits (*11*). The motifs show that decision signals entering BG across cortical-BG pathways can be selectively filtered by prefrontal circuits while signals output from BG can be subject to more widespread processing necessary to direct goal-oriented behavior. These results highlight a prominent role for β-oscillatory activity in modulating convergence and divergence of frontal-BG/Amyg communication within the mood pathways (*22*). Modulators in one network may, for example, also modulate the excitability of other networks and so may reveal mechanisms of interaction between the different networks integrating sensation, reward and action to guide motivated behavior. Such network mechanisms may underlie the search for novel biomarkers for mood disorders (*8*) and personalized treatment by using recordings to optimize spatiotemporal patterns of stimulation (*24, 25*).

Our empirical approach naturally complements mathematical models within the multilayer networks framework (*26, 27*) that investigate how functional brain networks reconfigure over time in order to predict the brain state associated with pathological functional connectivity. Taking a causal approach, however, offers important advantages. Functional interactions are often inferred based on the structure of correlations in neural activity and from examining responses to large-amplitude and/or stimulation pulse trains which are subject to several confounds. Correlations are sensitive to the confounding influence of common inputs from other brain regions, yielding network edges even when the receiver does not receive any input from the sender (*19*). Causal responses cannot be due to common input, which is consistent with their sparse presence in our data. Inferences from large-amplitude stimulation pulses or pulse trains suffer other confounds. Stimulating may recruit network responses due to other mechanisms (*28*), changing the interaction instead of probing it. Delivering isolated low-amplitude microstimulation pulses more directly probes network excitability while avoiding the confounding effects and network responses.

Prior studies have examined neural excitability by applying primary and secondary interventions at the sender only, perhaps reflecting technical limitations (*14, 29*). As a result, edge- and node-based mechanisms could not be dissociated. With sender-only interventions, changes in receiver node responses may reflect changes in pre-synaptic activity recruited by the primary intervention, or changes in the priming impact of the secondary intervention on the sender (*30*). Using our large-scale multisite approach, we disambiguated these effects and investigated dynamic changes in excitability across edges connecting different senders and receivers. Critically, we could do so independently of node activity at both the sender and receiver to specifically conclude that edge-based responses were disrupted.

The variable sender-receiver excitability we observed suggest orthodromic and trans-synaptic stimulation effects (*31–33*), and we cannot rule out antidromic stimulation effects. The causal network analysis we report, however, extends beyond putatively direct interactions to uncover mechanisms of neuromodulation that are not focal but distributed. The observed β-power suppression by gray matter SB-TetMS is consistent with the ‘synaptic filtering’ hypothesis. According to synaptic filtering, a high-frequency stimulation pulse train delivered to the pre-synaptic site induces short-term suppression that selectively suppresses the synaptic transmission of low-frequency content contained in stimulation-induced neural activity at the post-synaptic site (*34, 35*). Surprisingly, white matter SB-TetMS did not modulate receiver node activity overall. This was true even when receiver node responses to sender stimulation were suppressed. Our findings also reinforce prior work showing that high-frequency stimulation at white matter modulator sites can disrupt or block synaptic transmission from the sender to receiver (*4, 36*). Such effects are said to induce ‘information lesions’, highlighting the potential for SB-TetMS as a tool to suppress network excitability.

In conclusion, performing a minimally-perturbative causal network analysis reveals specific network mechanisms of multiregional communication and directly relates SB-TetMS neuromodulation to a general mechanism of network excitability that supports multiregional communication. These results reveal multiregional communication within the cortico-subcortical limbic mood processing network and point to how next-generation brain therapies can combine neuromodulation with neural decoding to correct disordered multiregional communication.

## Acknowledgements

We would like to thank Baldwin Goodell, Charles Gray, Jessica Kleinbart, and Amy Orsborn for assistance with chamber and microdrive system design; Stephen Frey and Brian Hynes for custom modifications to the Brainsight system; Keith Sanzenbach and Pablo Velasco from NYU Center for Imaging for help with magnetic resonance imaging and diffusion weighted imaging; Ryan Shewcraft, John Choi, Marsela Rubiano, Yoohee Jang, Octavia Martin, and NYU Office of Veterinary Resources for help with animal preparation and care. This work was supported, in part, by the Defense Advanced Research Projects Agency (DARPA) under Cooperative Agreement Number W911NF-14-2-0043, issued by the Army Research Office contracting office in support of DARPA’S SUBNETS program (to B.P.). The views, opinions, and/or findings expressed are those of the author(s) and should not be interpreted as representing the official views or policies of the Department of Defense or the U.S. Government. This work was also supported, in part, by an award from Simons Collaboration on the Global Brain (to B.P.), and US National Institutes of Health (NIH) BRAIN grant R01-NS104923 (to B.P.).

## Author Contributions

S.Q. and B.P. conceived and designed the experiment; S.Q., J.I.S., K.A.B., B.F. and B.P. performed the research; S.Q. and B.P. analyzed the data; S.Q. and B.P. wrote the manuscript. B.P. oversaw and guided all aspects of the project.

## Competing Financial Interests

There are no competing financial interests.

## MATERIALS AND METHODS

### Large-scale microdrive system design

We developed a customized large-scale semi-chronic microdrive system for mapping and manipulating the cortico-subcortical mood processing network. The screw-driven actuation mechanism of the microdrive provides bi-directionally independent control of the position of 220 microelectrodes (1.5-mm spacing) along a single axis with a range up to 32 mm (Monkey M) and 40 mm (Monkey A) with 125-µm pitch (*1*). Each actuator consists of a lead screw, a teardrop brass shuttle bonded to the electrode tail with conductive epoxy, and a compression spring. Each electrode can be moved with an accuracy of approximately 15 µm. For Monkey M, we distributed two types of microelectrode in the microdrive, 160 platinum/iridium (Pt/Ir) electrodes (MicroProbes, Gaithersburg, MD) with impedance 0.1–0.5 MΩ for extracellular recording and microstimulation, and 60 tungsten electrodes (Alpha Omega, Israel) for extracellular recording with impedance 0.8–1.2 MΩ. For Monkey A, we loaded 220 Pt/Ir electrodes (MicroProbes, Gaithersburg, MD) with impedance 0.5 MΩ for extracellular recording and microstimulation. Electrode impedances were measured at 1 kHz (Bak Electronics, Umatilla, FL). The tungsten electrode’s shank diameter was 125 µm and its total diameter was 250 µm with glass insulation. The Pt/Ir electrode’s shank diameter was 225 µm and its total diameter was 304 µm with parylene C and polyimide insulation.

### Magnetic resonance imaging and processing

We reconstructed each monkey’s brain, skull and cerebral vasculature (Brainsight^®^, Rogue Research, Montreal, QC) from anatomical magnetic resonance images (MRIs) and MRIs using ABLAVAR^®^ contrast agent for angiography with a T1-weighted magnetization-prepared rapid acquisition gradient-echo (MPRAGE) sequence. We also performed multishell high-angular resolution diffusion imaging (HARDI) tractography (*2*) registered to microdrive implantation. We collected diffusion weighted images to reconstruct white matter tracts connecting key cortical and subcortical areas of interest. During MRI procedures, the monkeys were anesthetized with isoflurane and placed in the scanner in sphinx position. We acquired data with a 3-Tesla (3T) Siemens Allegra (Erlangen, Germany) using 3 elements out of a 4-channel phased array from Nova Medical Inc. (Wilmington, MA) and 64 gradient directions in 1.2 mm^2^ in-plane resolution (TR = 7000 ms; TE = 110 ms; b-values: 0, 750, 1500, 2250 s/mm^2^; FOV: 80×64 pixels; slices: 48; slice thickness: 1.2 mm; DWI to *b*_0_ ratio 65:1). To correct for geometric distortions from field inhomogeneities caused by the non-zero off-resonance fields, we collected data with reversed phase-encode blips, forming pairs of images with distortions going in opposite directions. From these pairs, we estimated the susceptibility-induced off-resonance field using FSL’s TOPUP tool (*3*). We combined the two images into a single corrected image. We corrected Eddy currents generated by the 64 gradient directions using FSL’s eddy tool (*4*).

### Experimental preparation

All surgical and experimental procedures were performed in compliance with the National Institute of Health Guide for Care and Use of Laboratory Animals and were approved by the New York University Institutional Animal Care and Use Committee. Two male rhesus macaques (*Macaca mulatta*) participated in the study (Monkey M, 8.4 kg and Monkey A, 7 kg at the beginning of the experiments). We performed a craniotomy and dura thinning on the targeted brain regions over the left (Monkey M) and right (Monkey A) hemispheres. We implanted a customized large-scale recording chamber (Gray Matter Research, Bozeman, MT) fitted to the skull surface using MR-guided stereotaxic surgical techniques (Brainsight^®^, Rogue Research, Montreal, QC). We aligned the chamber and registered it within 1 mm of the target coordinates (nominally 0.4 mm) and affixed and sealed it to the skull surface via C&B-METABOND^®^ (Parknell Inc., Edgewood, NY) and dental acrylic. We then mounted the microdrive into the chamber and sealed it with compressed gaskets and room-temperature-vulcanizing (RTV) sealant (734 flowable sealant, Dow Corning, Midland, MI). To target each brain region, we registered electrodes to anatomical magnetic resonance images (MRIs) and magnetic resonance angiograms (MRAs). To limit routes for infection, we customized the shape of the chamber and microdrive to each animal’s anatomy and established that the chamber was sealed following implantation. To limit damage to the vasculature, midline, and ventricles when lowering electrodes, we only advanced a subset of electrodes along trajectories considered safe, greater than 2 mm from MR-visible vasculature, ventricles, and midline. Finally, we co-registered the MRIs to the MNI Paxinos labels (*5*) to label each recording and stimulation site (**fig. S1**). We successfully studied activity from 165 electrodes (50 tungsten and 115 Pt/Ir) in Monkey M for over 24 months and 208 Pt/Ir electrodes in Monkey A for over 12 months.

We tested long-term stability of neural recordings in two ways. First, we examined local field potential (LFP) activity during the movement epoch of a reaching task. **Figure S1A-C** presents task-evoked LFP activity for 67 depths along a sample electrode trajectory. Task-evoked LFP activity clearly increased and decreased in magnitude as the electrode transitioned into and out of gray matter, respectively. We also validated localization based on spiking. Increased task-evoked LFP activity was associated with increasing spiking activity (**fig. S1, D-G**). Patterns of spiking activity and task-evoked LFP activity were preserved over many months, demonstrating long-term stability of the implanted device.

### Estimation of sampled brain network nodes and edges

We consider each brain region to be a network node. We sampled each node by positioning an electrode at a site in the node. For every pair of electrodes that simultaneously recorded activity in two nodes, we obtained one sample of the network edge connecting the two nodes. Repeated node sampling enabled repeated edge sampling. With M electrodes positioned at different sites in Node1, and N less than M electrodes in Node2, we obtained N non-overlapping samples of the Node1-Node2 network edge. In Monkey M, we sampled four cortical nodes at 3521 sites (1506 OFC, 890 ACC, 817 dPFC, and 308 M1 sites) and three subcortical nodes at 1125 sites (915 CN, 122 Put, and 88 GP sites). In Monkey A, we sampled seven cortical nodes at 3900 sites (319 OFC, 469 ACC, 343 PCC, 427 dPFC, 713 M1, 429 S1, and 1200 PPC sites) and eight subcortical nodes at 997 sites (614 CN, 214 Put, 59 GP, 45 Amyg, 25 PrS, 20 SI, 2 BFB, and 18 NAc). We targeted our causal network analysis to 9548 samples from the cortico-subcortical limbic network across two monkeys, specifically for 1359 OFC•ACC, 1244 OFC-dPFC, 1529 OFC-CN, 336 OFC-Put, 45 OFC-Amyg, 1244 ACC-dPFC, 1359 ACC-CN, 336 ACC-Put, 45 ACC-Amyg, 1244 dPFC-CN, 336 dPFC-Put, 45 dPFC-Amyg, 336 CN-Put, 45 CN-Amyg, and 45 Put-Amgy interactions. Note that OFC combines medial OFC and lateral OFC; dPFC combines inferior, middle, and superior frontal gyri; and posterior parietal cortex (PPC) combining superior parietal lobule (SPL), supramarginal gyrus (SMG), and precuneus (pCun).

### Neural recordings and microstimulation

The monkeys were awake, head-restrained and seated in a primate chair placed in an unlit sound-attenuated electromagnetically shielded booth (ETS Lindgren). Neural recordings were referenced to a ground screw implanted in the left posterior parietal lobe (Monkey M) and left occipital lobe (Monkey A), respectively, with the tip of the screw just piercing through the dura mater. Neural signals from all channels were simultaneously amplified and digitized at 30 kHz with 16 bits of resolution with the lowest significant bit equal to 0.1 μV (NSpike, Harvard Instrumentation Lab; unit gain headstage, Blackrock Microsystems), and continuously streamed to disk during the experiment with lights switched off in the recording booth.

We applied microstimulation using a bipolar configuration, made by simultaneously sending a biphasic charge-balanced square wave pulse via a pair of Pt/Ir microelectrodes with the same pulse amplitude, pulse width, and interphase interval, but opposite polarity (e.g. cathode-lead for electrode 1 and anode-lead for electrode 2) (Cerestim R96, Blackrock Microsystems, Salt Lake City, UT). We used the pulse width as 100 µs per phase and interphase interval as 53 µs for all stimulation sessions. The Pt/Ir microelectrodes had a typical tip geometric surface area of 223 ± 37 µm^2^. For instance, a single pulse, with amplitude of 40 µA and width of 100 µs per phase (4 nC/ph), could yield a charge density of approximately 1800 µC/cm^2^. We simultaneously recorded neural signals from all electrodes while stimulating at certain pair of electrodes. No seizure activity due to stimulation was detected during experiments.

We implemented two microstimulation protocols to perform causal network analysis. Primarily used in this study, we designed a novel multisite spatiotemporal patterned microstimulation framework to assess changes in the weight of network edges identified using evoked LFP responses to single microstimulation pulses. For each stimulation bout, the stimulation pattern started with a Pre-tetanic single microstimulation pulse from a pair of electrodes at a network site (sender) to identify the network edges, followed by a TetMS from another pair of electrodes at another network site (TetMS-modulator), and followed by a Post-tetanic single microstimulation pulse at the sender. The latency between the Pre-tetanic pulse and the onset of the TetMS (*τ*_Pre_) was either 0.5 or 1 s. The duration (*T*) of the TetMS pulse train and the latency (*τ*_Probe_) between the offset of TetMS pulse train and the onset of Post-tetanic single pulse varied in a pseudo-random fashion bout by bout (T: range = 50-2000 ms; T_post_: range = 0-200 ms; mean: 57 ms; and median: 50 ms; **table S1**). We also varied the inter-trial interval bout by bout (duration of each stimulation bout plus 2-4 s variation). We summarized all stimulation parameters in **table S1**. We defined the neural excitability (edge weight) of identified network edges as the peak magnitude of pulse-triggered averaged evoked LFP response within a 100-ms post single-pulse epoch. We also used a single-pulse microstimulation protocol with either periodic or Poisson pulse train (*6*) to identify network edges. This protocol consisted of 1-s pulse trains and 1-3 s baseline epochs. Due to stimulation artifact, to preserve a minimum 100-ms post single-pulse epoch for analyzing the response to each of pulses, we used pulse train frequencies of 5 and 10 Hz with corresponding refractory period of 200 and 100 ms, respectively.

Of 266 stimulation site-amplitude combinations tested to establish the threshold microstimulation, 16 site-amplitude combinations revealed 78 significant functional interactions between the sender and receiver in 21 network edges. Bipolar microstimulation limited the current spread from the stimulation site and corresponding stimulation artifacts. Bipolar re-referencing for recordings aimed to reveal focal evoked responses and also limited stimulation artifacts. The secondary intervention was designed to be minimally-perturbative. By stimulating the sender above threshold and recording the receiver at threshold, the receiver response to input from the sender measured the weight of the network edge (sender→receiver). To do so, we stimulated the sender with different current amplitudes. We then set the stimulation dose at the threshold needed to generate a stimulation-evoked response at receiver sites. The established threshold for single pulses ranged from 10 to 100 µA with a median amplitude of 40 µA (**table S1**).

### Pulse-triggered evoked potentials

We obtained LFP activity offline by low-pass filtering the broadband raw recording at 400 Hz using a multitaper filter with time duration of 0.025 s, frequency bandwidth of 400 Hz, and center frequency of 0 Hz, and then down-sampled to 1 kHz from 30 kHz. To limit the common-mode confounds of the neural recordings, we digitally referenced the neural signal of each channel to its nearest neighbor within 3 mm based on the electrode depth. We removed the events from the analysis if they exceeded 10 standard deviations of from the mean across the stimulation event pool. We also removed noisy or bad channels via visual inspection. Pulse-triggered evoked potentials were computed by averaging LFP signals aligned to the onset time of each single pulse. The z-scored pulse-triggered evoked potentials were then computed using the standard deviation (SD) of the baseline (-30 to -5 ms). We computed the baseline SD for each electrode separately using all time points in the baseline window of the pulse-triggered evoked potential.

We should note that suppression of the evoked response to the Post-tetanic single pulse we observed may be confounded by an effect related to the phase of inhibition after the excitatory response to the Pre-tetanic single pulse (*7–9*) or related to suppressed responses to paired microstimulation pulses (*9, 10*). Our experimental design employed Pre-tetanic and Post-tetanic pulses separated by 0.75-3 s, mitigating such concerns.

### Pulse-triggered multiunit activity

We obtained multiunit activity (MUA) offline by first band-pass filtering the broadband raw recording from 0.3 to 6.6 kHz with time duration of 0.01 s, frequency bandwidth of 3 kHz, and center frequency of 3.3 kHz. We then applied a 3.5 standard deviation threshold to identify putative spikes (1.6-ms duration). We labeled all waveforms that exceeded this peak as multiunit action potentials. We computed multiunit peristimulus time histograms (PSTH) by aligning MUA to the onset time of each single pulse. Individual spikes were collected in 1 ms bins and the corresponding histogram was smoothed by convolving with a Gaussian function with a standard deviation of 5 ms. Data from -1 ms to 5 (or 10) ms around the single pulse onset were not used for multiunit activity analysis due to stimulation artifact.

### Detection of pulse-evoked LFP responses using stimAccLLR model

To quantify the detection of evoked responses to microstimulation on a single-pulse basis, we used the stimulation-based accumulating log-likelihood ratio (stimAccLLR) method to determine when selectivity in the LFP activity for the null and alternative hypotheses emerged, i.e. the latency of a stimulation response. The latency from single microstimulation pulses was determined by the time at which Pulse Event reached a detection threshold. The threshold was selected based on a trade-off between speed (latency) and accuracy (probability of correct classification) as we increased the level of the detection thresholds from AccLLR equal to zero (fig. S4).

In the stimAccLLR method, we defined a probabilistic model of the LFP activity for the two alternatives being tested. To determine the latency for a response to single microstimulation pulses, we define the two alternative hypotheses into “condition 1” and “condition 2”, where “condition 1” represents the LFP activity (Pulse Event) in the post-stimulus epoch, while “condition 2” represents LFP activity (Pulse Null) in the pre-stimulus epoch. We performed stimAccLLR on all stimulation response to single microstimulation pulses, combining all Pre-tetanic single pulses using the multisite spatiotemporal patterned microstimulation protocol and all the single pulses using the Poisson burst microstimulation protocol. For Pre-tetanic single pulses, Pulse Event was the 100-ms LFP activity after Pre-tetanic pulse onset plus 5 or 10-ms blanking epoch due to stimulation artifact, while Pulse Null was the 100-ms LFP activity before Pre-tetanic pulse onset. For single pulses in the Poisson burst, Pulse Event was the 50-ms LFP activity after each pulse onset plus 5 or 10-ms blanking epoch due to stimulation artifact, while Pulse Null was the 50-ms LFP activity before each pulse onset.

We modeled LFP activity as independent observations from an underlying Gaussian distribution. The signal *x*(*t*) at time *t* is expressed as a mean waveform *µ*(*t*) with additive Gaussian noise, *ε*(*t*)

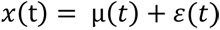

The likelihood of observed data *x*(*t*) being generated by each model was given by

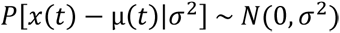

The likelihood ratio, *LLR*(*t*), at time *t* for two time-varying Gaussian LFP models, assuming the noise in both models had the same variance, *σ*^2^, was given by

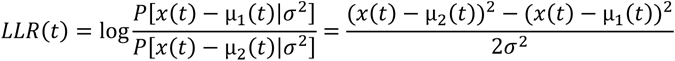

Finally, we calculated the accumulated log-likelihood ratio by summing log-likelihoods over time:

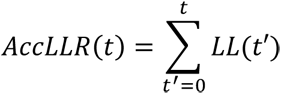

To quantify signal selectivity, we performed a receiver-operating characteristic (ROC) analysis on the AccLLR values. Analysis was performed every millisecond to discriminate activity from “condition 1” and “condition 2” events. We defined overall signal selectivity as the choice probability from the ROC analysis at the end of a 100-ms or 50-ms accumulation interval. We summarized the detectable likelihood of the stimulation response to a single microstimulation pulses between brain regions. We run a binomial test at the significant level of 0.05 to determine if some causally sampled network edges that showed significant directed functional interactions were more likely selective than others.

Performing stimAccLLR reveals ‘Hit’ and ‘Miss’ events in the receiver responses. To further test for the presence of two response classes, we classified pulse-by-pulse evoked responses using a Gaussian mixture model (GMM). The GMM clustered pulse-evoked responses into two classes. For a majority of the identified network edge samples (50/78, 64%) the analysis methods agreed (*P* < 0.05, Fisher’s exact test, two-tailed test; **fig. S8**).

### Edge- and node-modulation analysis

With the multisite spatiotemporal patterned microstimulation framework, we observed suppression of neural excitability of all identified edge samples after applying short-burst tetanic microstimulation (SB-TetMS). It is worth noting that, instead of only observing suppressed network excitability, we may also see both increased and decreased excitability at longer time scales by altering bidirectional synaptic plasticity when using, for instance, phase-dependent stimulation for the primary intervention (*11*). For analyzing receiver node activity before and after SB-TetMS. We focused the analysis on LFP activity because it is clinically-important and yielded reliable, long-term recordings across the sampled network. We quantified node activity during 100 ms immediately before (Pre-state) and 150-250 ms after (Post-state) the pulse-SB-TetMS-pulse sequence using root mean square (RMS) power in beta (β: 13-30 Hz) frequency band, a key signature of node activity used in DBS studies (*12*), and in high-gamma (high-γ: 70-150 Hz) frequency band, a key signature of node activity that reflects local spiking of populations of neurons (*13*). Since receiver response to the Post-tetanic pulse is due to the sender, we defined the Post-state as activity shortly after the pulse-triggered evoked response.

### Identifying modulators and decoding sender-receiver communication

To identify modulators for each of identified network edges, we grouped the Pre-tetanic pulse baseline LFP activity (500-ms epoch before pulse onset) of all recorded sites into two categories based on ‘Hit’ and ‘Miss’ events detected from the stimAccLLR procedure. We estimated the power spectral density (PSD) of pre-pulse baseline LFP activity using multitaper methods with 500-ms sliding window with ±2-Hz smoothing. We removed the events from the analysis if they exceeded 5 standard deviations of from the mean across the stimulation event pool. We then performed the ROC analysis to test if the respective LFP power of each recorded site in ‘Hit’ events was significantly different from ‘Miss’ events in the frequency band of 10-40 Hz. For a given identified network edge, we defined the recording site in the gray matter with the area under ROC curve (AUC) greater than a threshold as modulators that significantly modulated receiver responses, where the threshold was set when AUC was 95% confidence interval above chance (two-tailed test, FDR-corrected *P* < 0.05). We focused on this analysis using the data collected in the multisite spatiotemporal patterned microstimulation framework. From 62 identified sender-receiver pairs, we identified 63 modulator sites (1797 sites tested) from 20 samples across 11 network edges (**table S3–S5**).

### Spectrogram of modulator baseline activity

To show how the baseline neural activity of tested modulators significantly modulated sender-receiver communication, we first sorted the events by detected ‘Hit’ and ‘Miss’ events at the receiver. We then estimated spectrograms of modulator baseline LFP activity (1-s epoch before Pre-tetanic pulse onset) in ‘Hit’ and ‘Miss’ events, respectively, using multitaper methods with 500-ms sliding window with ±5-Hz smoothing and 5-ms stepping between spectral estimates. We tested the difference in LFP power between ‘Hit’ and ‘Miss’ events against a null hypothesis that there was no LFP power difference using a permutation test (10,000 permutations). To generate the null distribution for no LFP power difference, we randomly shuffled the order of combined ‘Hit’ and ‘Miss’ events, followed by computing shuffled LFP power difference between ‘Hit’ and ‘Miss’ events. For the significant regions presented in the spectrograms (*P* < 0.05, two-tailed test), we applied a cluster correction to correct for multiple comparisons at the significant level of 0.05 (*14*).

## SUPPLEMENTARY TEXT

### Minimally-perturbative causal network analysis

Our inferences depend on using a minimally-perturbative secondary intervention, for which we present multiple lines of supporting evidence. First, we used single, bipolar, biphasic pulses of relatively low-amplitude that are not necessarily expected to generate large-scale network responses. Indeed, single microstimulation pulse has not been previously used to map large-scale networks in the awake primate brain. The natural concern is that network responses following such minimal perturbations may not be detectable. Second, following single microstimulation pulses we observed sparse directed functional interactions instead of widespread network effects. Approximately 0.6% of sampled sites showed statistically-significant responses. We should note that sparsity may result from the presence of residual stimulation artifacts that obscure responses at some sites. The degree of sparsity likely also reflects the minimally-perturbative microstimulation approach we used, which is conservative and so may fail to drive detectable responses at some sites where more pre-synaptic inputs might be required to produce excitatory post-synaptic activity (*15*). Sparsity also likely reflects the underlying anatomical connections and the fact that many recording sites may not receive inputs from the sender. Third, although the responses were sparse, they were strong, visible on a single-pulse basis, and varied moment-by-moment. The presence of strong pulse-evoked responses demonstrates the efficacy of the intervention and mitigates concern about false positives due to multiple comparisons involving the number of tested network edges. The absence of pulse-evoked responses at other times also demonstrates the receiver is perturbed at threshold and hence that the secondary intervention is unlikely to be altering interactions across the network.

### Alternative interpretations of changes in excitability

Changes in excitability likely reflect additional mechanisms within the receiver node. 1) SB-TetMS may change the excitatory-inhibitory balance in the receiver (*16*), reducing excitatory and increasing inhibitory potentials, consistent with reductions in firing rate as well as the pulse-evoked LFP responses we observed; 2) SB-TetMS may also induce spatial reorganization of the pre-synaptic inputs that changes the strength of effective dipole moment generating the observed LFP activity (*17*); 3) The absence of responses following SB-TetMS that we observed might also reflect short-term depression in the post-synaptic neurons; 4) Momentary failures at the pre- and post-synaptic junction may also contribute to the gating of sender synaptic input to the receiver. Nevertheless, the predictability of neural excitability by modulators cannot simply be explained by mechanisms within the receiver node. Modulator decoding results provide strong evidence for network excitability-based mechanism for communication.

**Fig. S1.**
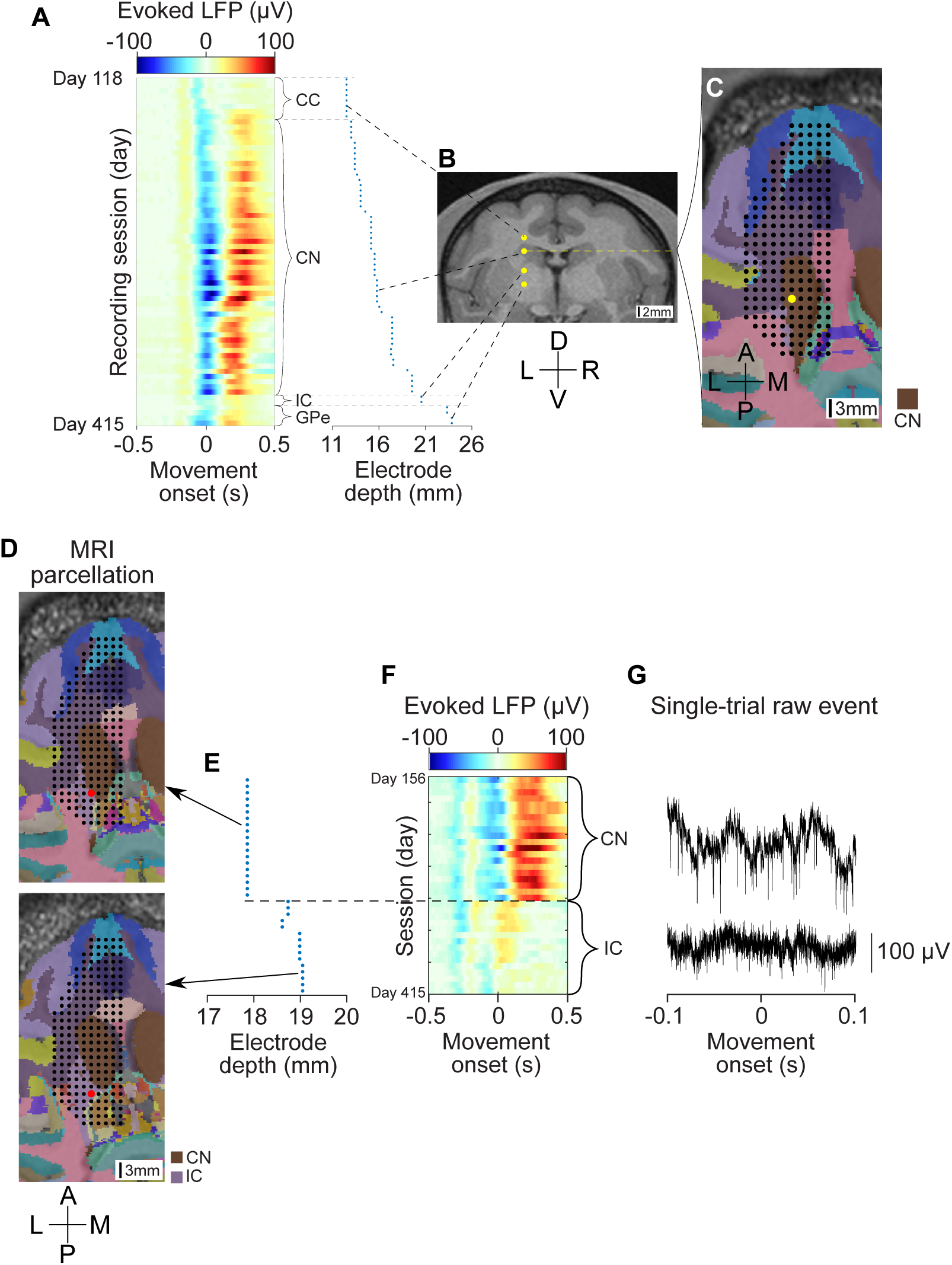
Validating anatomical targeting using depth profile of task-evoked LFP activity patterns. (**A**) White and gray matter transitions of average evoked LFP patterns recorded from an exmaple electrode in reach movement epoch as the electrode is advanced through corpus callosum (CC), caudate nucleus (CN), internal capsule (IC), and globus pallidus external (GPe). Electrode depth from cortical surface. (**B**) Co-registered four samples of the electrode depth in CC, CN, IC, and GPe on the coronal MRI slice. (**C**) Co-registered electrode tracks with anatomical parcellation of frontal lobe of Monkey M. Black dots: overlaid electrode grid. Yellow dots: the exmaple electrode at the horizontal MRI slice depth in CN. (**D-G**) Co-registrated electrode track of another example electrode at two different horizontal MRI slice depths (top: CN; bottom: IC). White and gray matter transitions of average evoked LFP patterns recorded from the exmaple electrode in reach movement epoch as the electrode is advanced through CN and IC. Higher evoked LFP activity in CN correlates with higher spiking activity. Anterior (A), posterior (P), left (L), right (R), medial (M), lateral (L), dorsal (D), and ventral (V) directions shown.

**Fig. S2.**
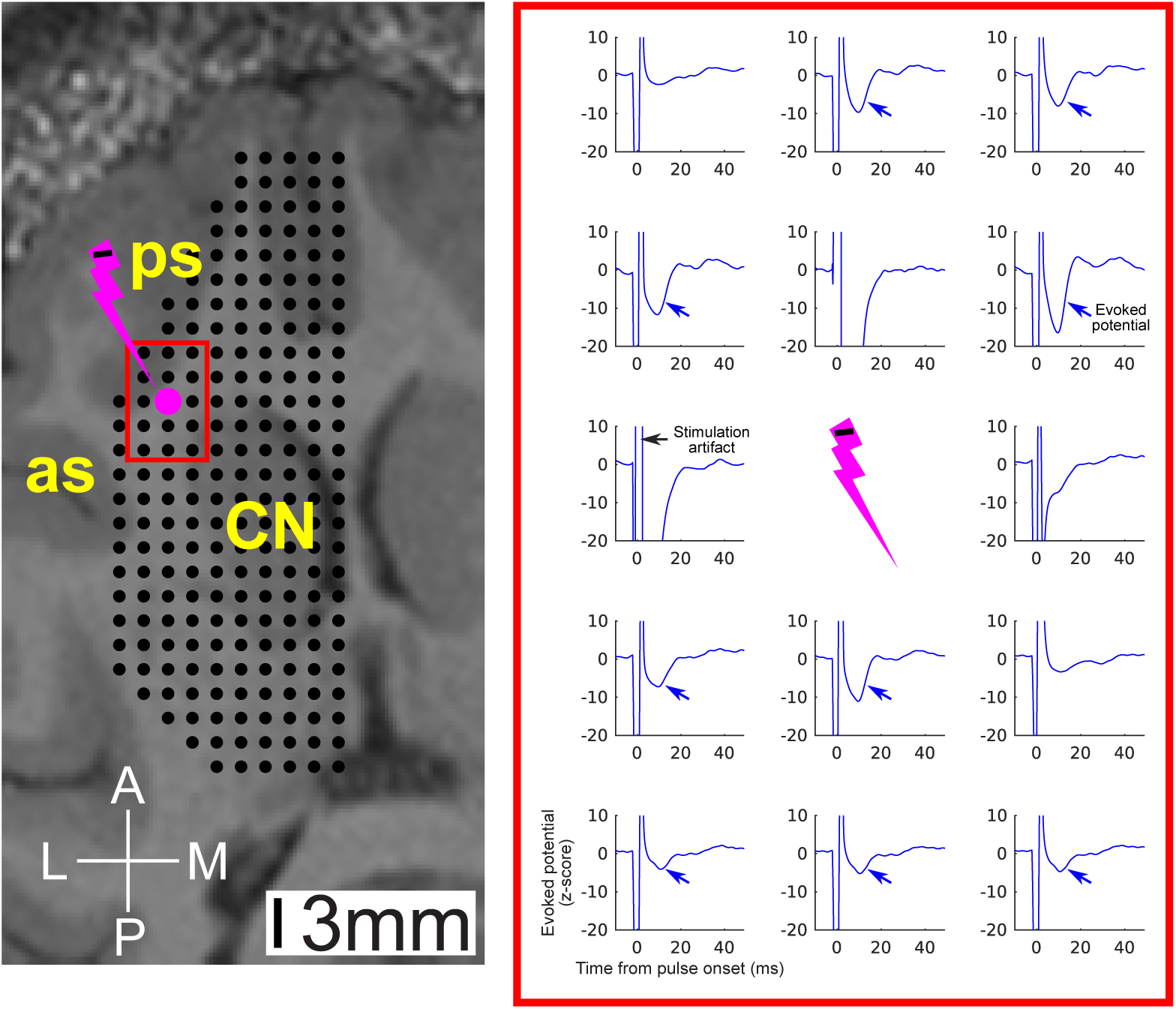
Example evoked responses to monopolar sinlge-pulse microstimulation. Pulse-triggered evoked potentials due to single, monopolar, biphasic, charge-balanced cathode-lead microstimulation pulses (40 μA, 100 μs/ph), demonstrating stimulation responses at the near-stimulation sites might be contaminated by stimulation current spread. CN: caudate nucleus; as: arcuate sulcus; ps: principal sulcus. Anterior (A), posterior (P), lateral (L), and medial (M) directions shown.

**Fig. S3.**
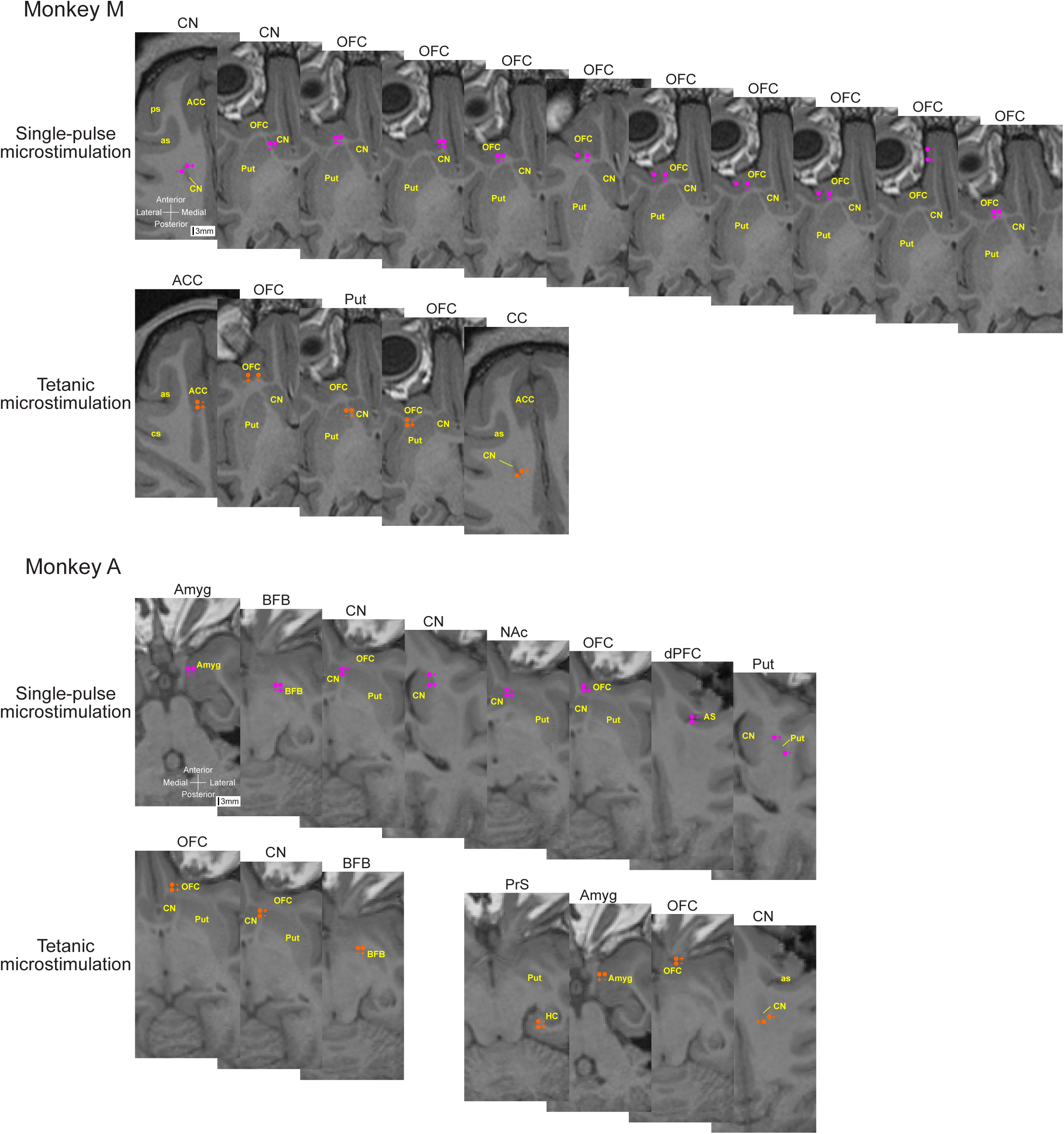
Anatomical locations of the single-pulse and tetanic bipolar microstimulation sites overlaid with horizontal MRI slices. Negative sign indicates cathode and positive sign indicates anode.

**Fig. S4.**
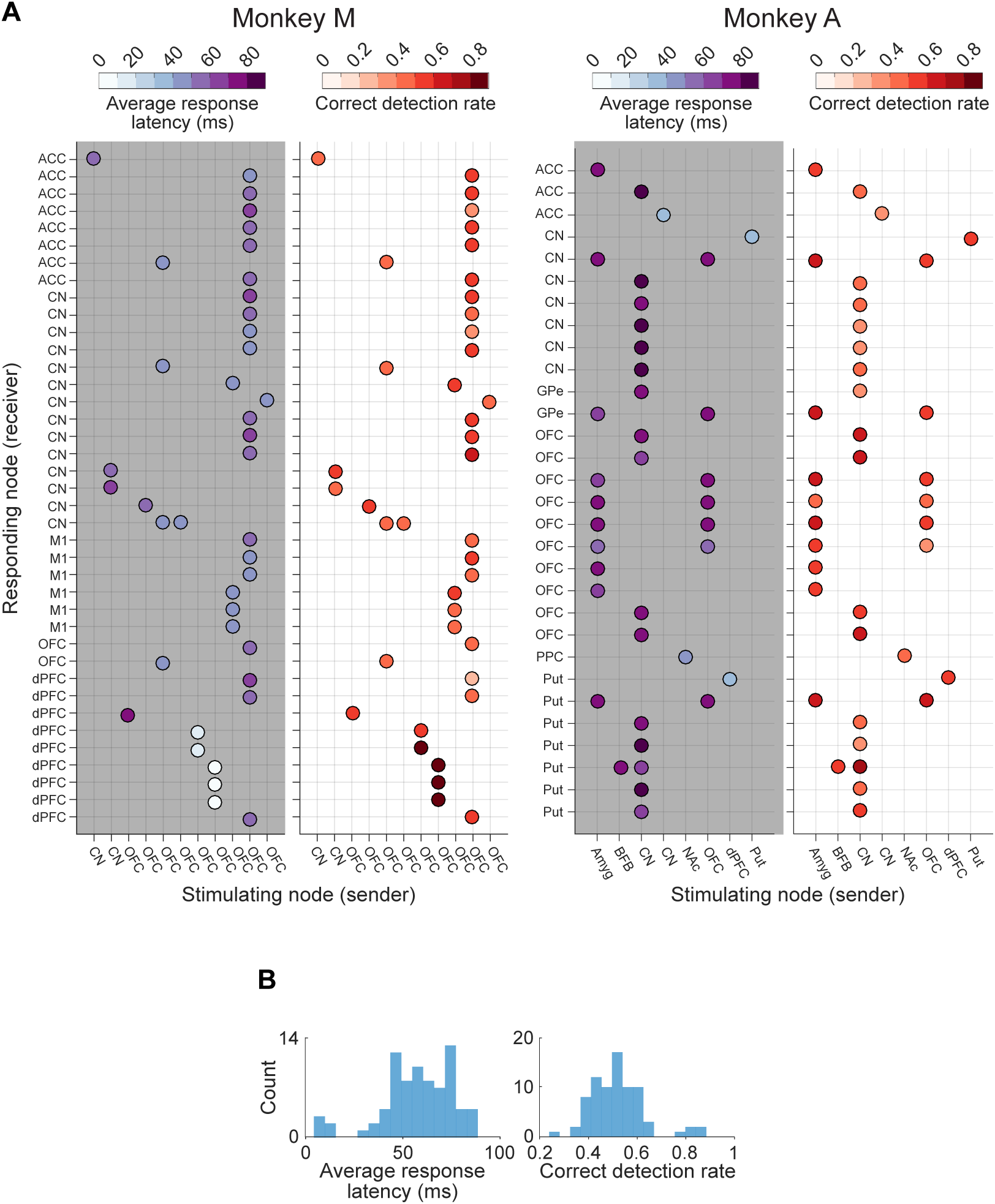
Population statistics of identified sender-receiver communication. (**A**) Average response latencies and correct detection rates of sender-receiver communication (Monkey M: 40; Monkey A: 38) (**B**) Population histogram of average response latencies and correct detection rates.

**Fig. S5.**
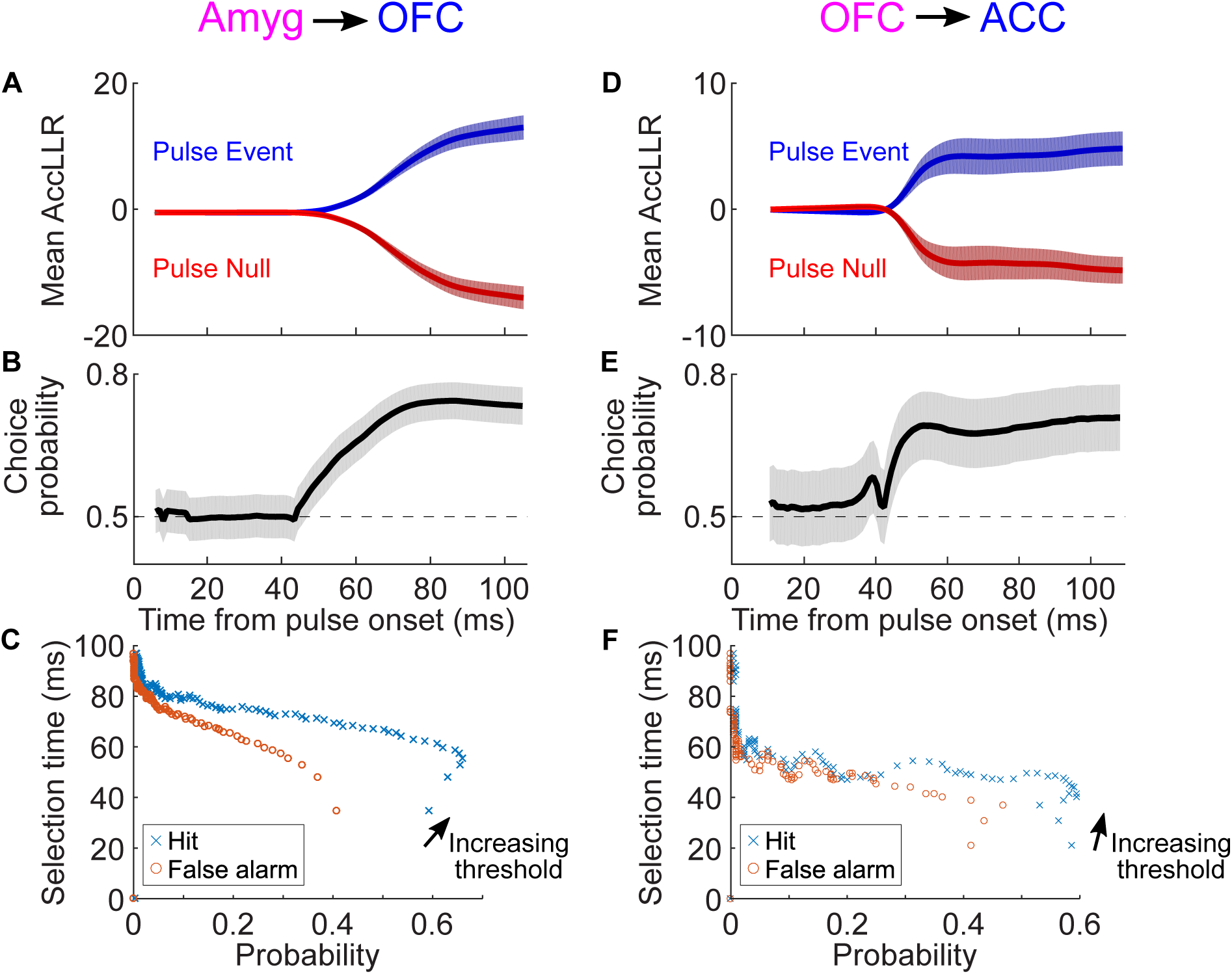
Identification of network edges using stimAccLLR. (**A-C**) The stimAccLLR results for the example Amyg→OFC edge in Monkey A. (**A**) Mean AccLLR of all stimulation trials during the pre-stimulus (red, Pulse Null) and post-stimulus (blue, Pulse Event) epoch (100-ms window). Error values are s.e.m., *n* = 322. First 5-ms data after the stimulation onset were not used due to stimulation artifact. (**B**) Choice probability from the receiver-operating characteristic (ROC) analysis applied to AccLLR traces at each time bin following onset for null and event. The shaded area represents 95% confidence intervals. (**C**) Probability of correctly detecting a single pulse from Pulse Event, ‘hit’ (cross), and incorrectly detecting a single pulse from Pulse Null, ‘false alarm’ (circle), plotted against the selection time as the level of the detection threshold is varied. The best correct detect performance decoding single microstimulation pulse of receiver response was 64%, with a 30% false alarm and 6% unknown. The average response latency at this false alarm rate was 64 ms. (**D-F**) The stimAccLLR results for the example OFC→ACC edge in Monkey M (*n* = 110). The best correct detect performance decoding single microstimulation pulse of receiver response was 50%, with a 25% false alarm and 25% unknown. The average response latency at this false alarm rate was 48 ms.

**Fig. S6.**
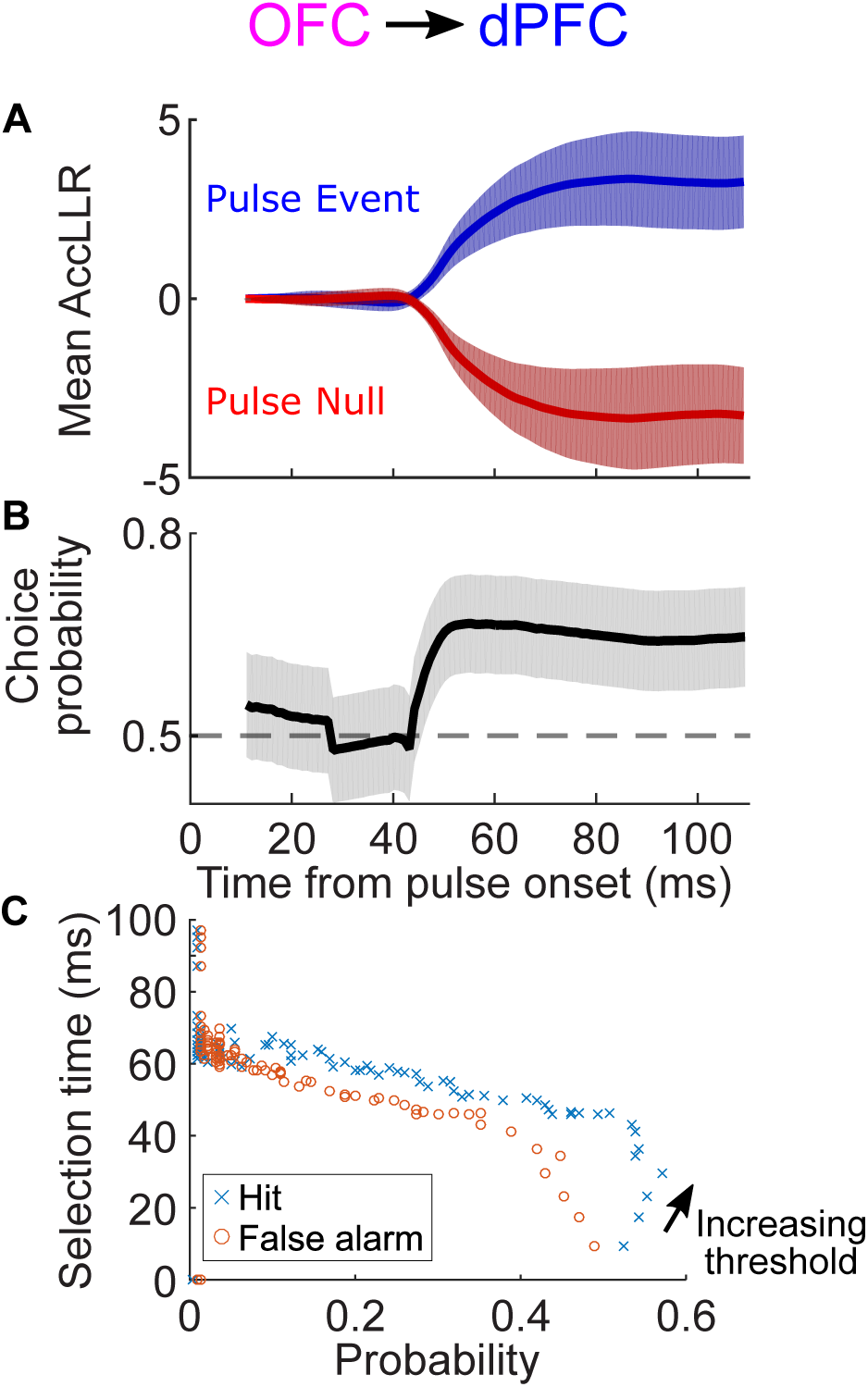
Identification of the example network edge OFC→dPFC using stimAccLLR. (**A**) Mean AccLLR of all stimulation bouts during the pre-stimulus (red, Pulse Null) and post-stimulus (blue, Pulse Event) epoch (100-ms window). Error values are s.e.m., *n* = 110. First 10-ms data after the stimulation onset were not used due to stimulation artifact. (**B**) Choice probability from the receiver-operating characteristic (ROC) analysis applied to AccLLR traces at each time bin following onset for null and event. The shaded area represents 95% confidence intervals. (**C**) Probability of correctly detecting a single pulse from Pulse Event, ‘hit’ (cross), and incorrectly detecting a single pulse from Pulse Null, ‘false alarm’ (circle), plotted against the selection time as the level of the detection threshold is varied.

**Fig. S7.**
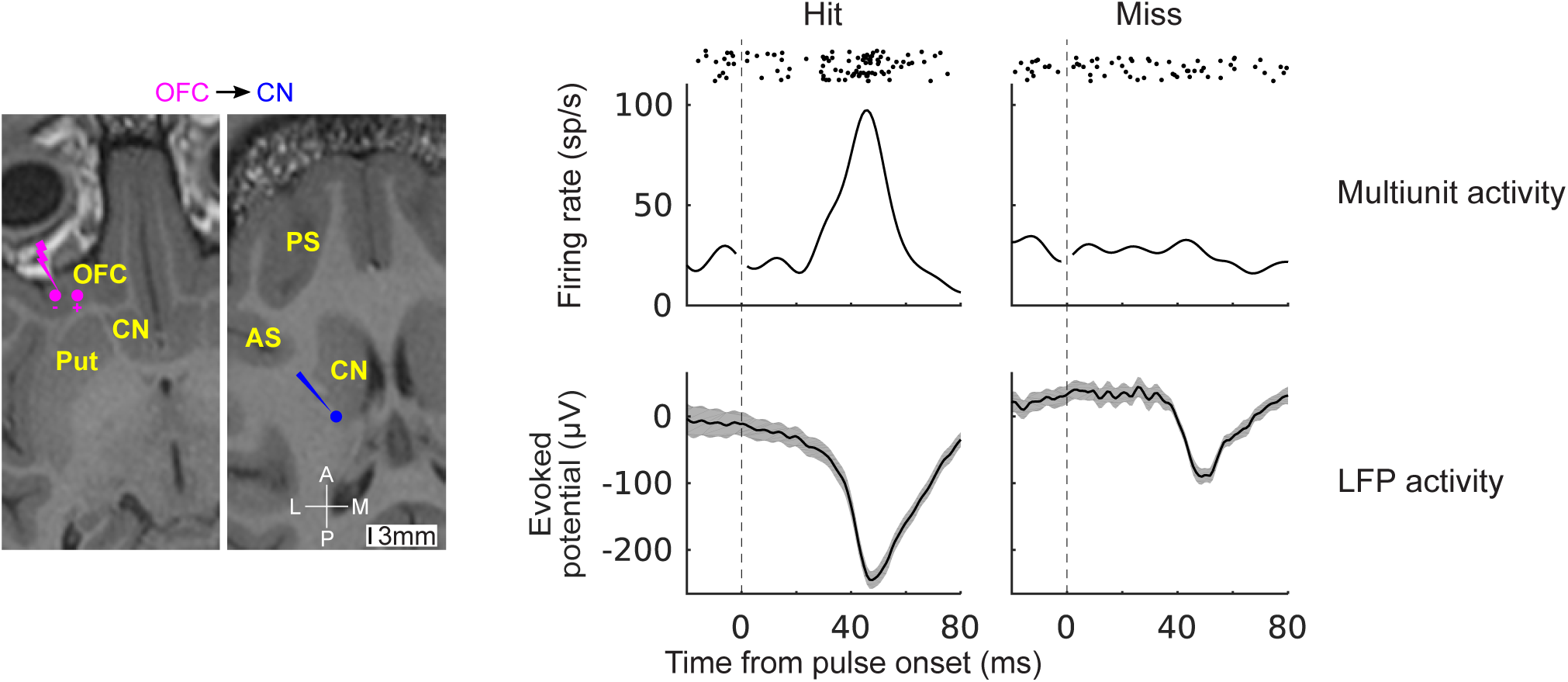
Driven multiunit activity and LFP activity of the caudate receiver in ‘Hit’ and ‘Miss’ events. Neural activity recorded in a CN receiver was driven by single, bipolar, biphasic, charge-balanced cathode-lead microstimulation pulses (10 μA, 100 μs/ph) delivered at an OFC sender. Driven multiunit activity are shown as raster plots and peristimulus histograms (PSTH). Shaded: +/-1 s.e.m. Anterior (A), posterior (P), lateral (L), and medial (M) directions shown.

**Fig. S7.**
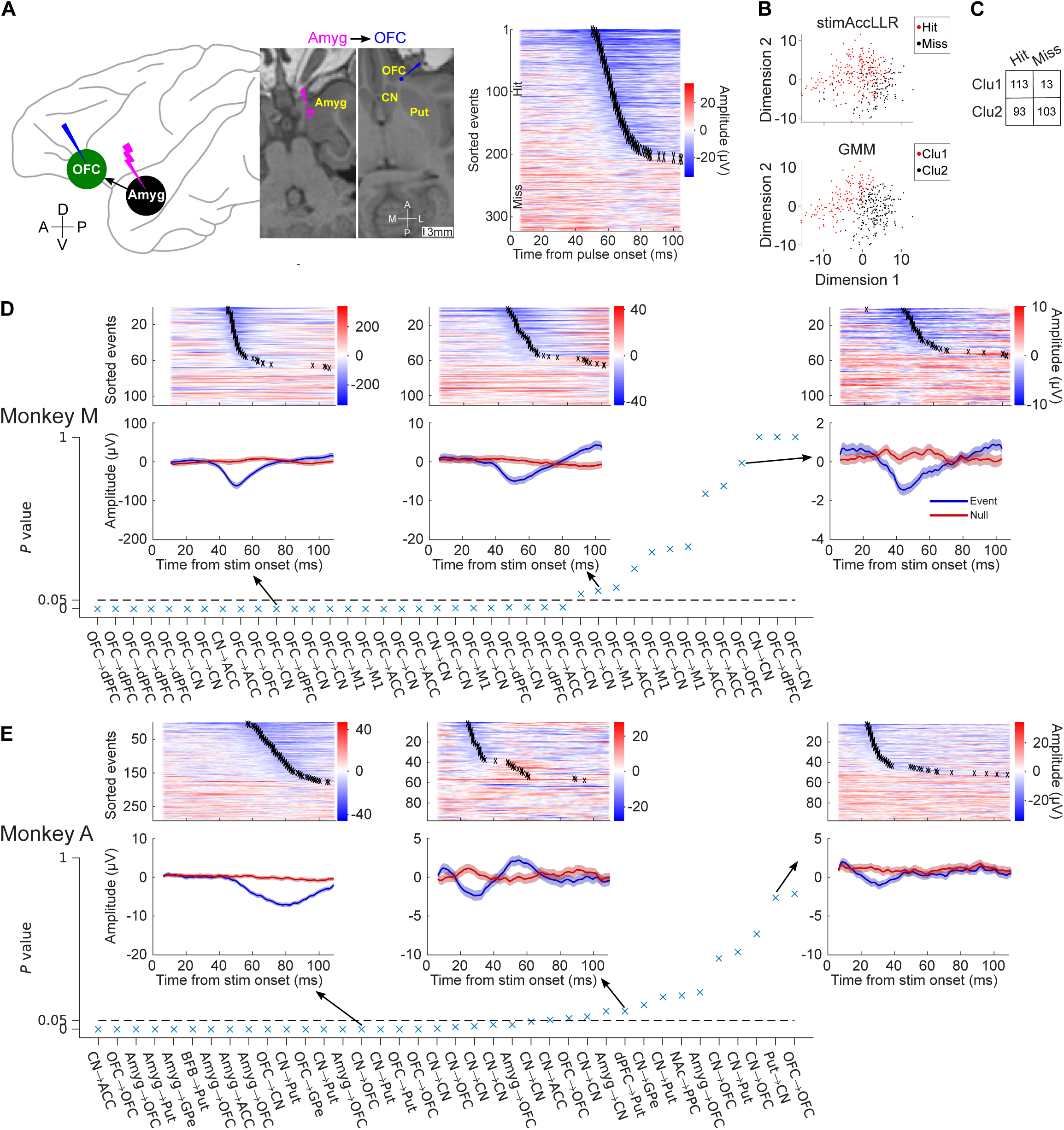
Fisher’s exact test for stimAccLLR and GMM classification results. (A) An example causally sampled network edge Amyg→OFC with the heat map of the pulse-by-pulse stimulation responses sorted by latencies. Anterior (A), posterior (P), lateral (L), and medial (M) directions shown. (**B**) Cluster plots of the waveforms of Pulse Events (100-ms epoch, 5 ms after pulse onset) between the first two projected dimensions. Left: color-coded based on the stimAccLLR result showing correctly ‘Hit’ vs ‘Miss’ events; Right: color-coded using the Gaussian mixture model (GMM) classifer. (**C**) Contingency table for Fisher’s exact test on the data shown in (B). *P*-value of 9.05e-16 indicates that there is significant association between the classification results using AccLLR and GMM (two-tailed test, significance level = 0.05). (**D**-**E**) P-values of all causally sampled network edges that showed directed funtional interactions and example stimAccLLR results.

**Table S1.**
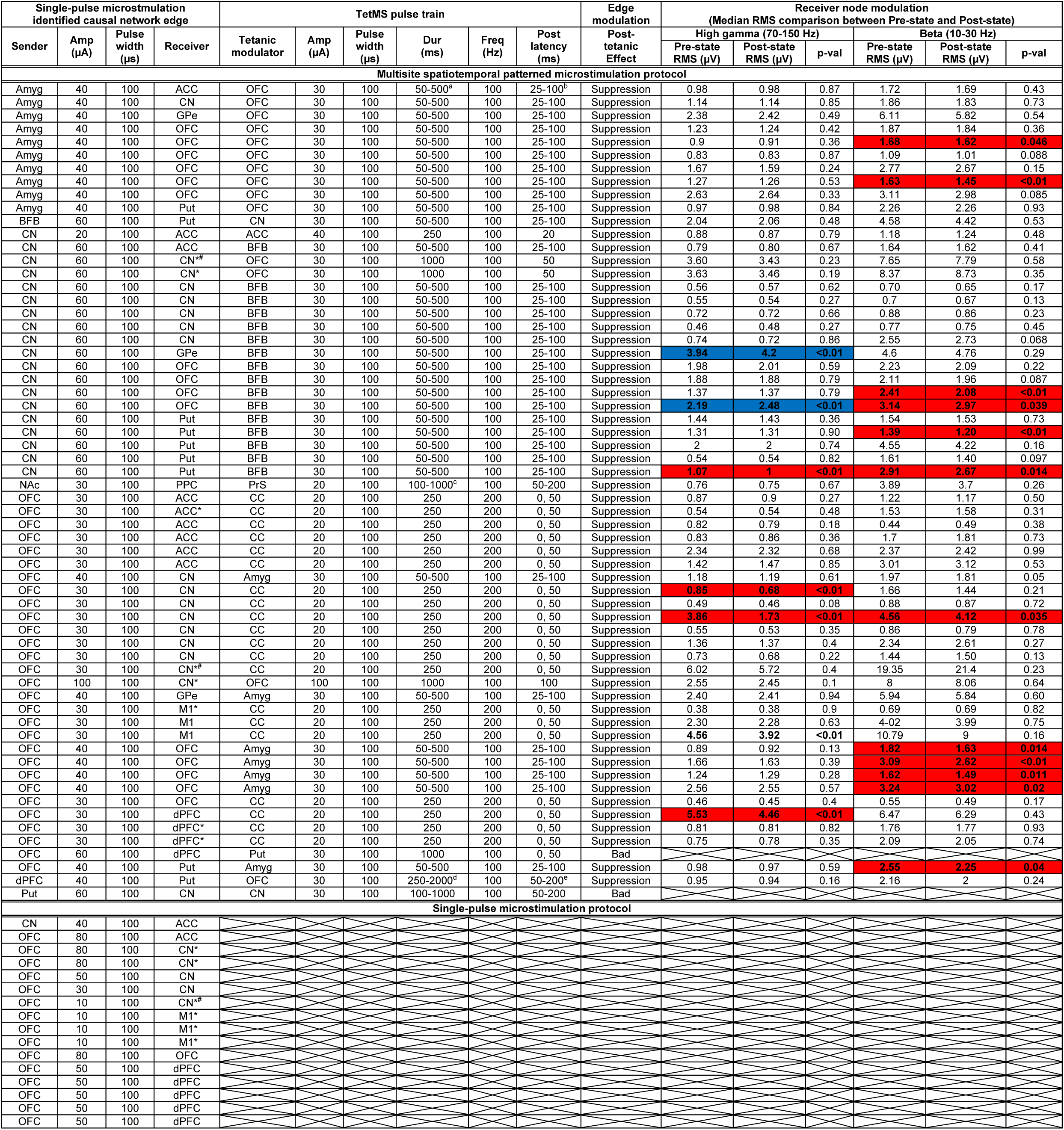
Summary of stimulation parameters for 78 directed network edge samples.

Notes:

<u>Cortical regions</u> ― OFC: orbitofrontal cortex; ACC: anterior cingulate cortex; dPFC: dorsal prefrontal cortex; M1: primary motor cortex; PPC: posterior parietal cortex

<u>Subcortical regions</u> ― CN: caudate nucleus; Put: putamen; GPe: globus pallidus external; Amyg: amygdala; BFB: basal forebrain; NAc: nucleus accumbens; PrS: presubiculum

<u>White matter</u> ― CC: corpus callosum

^a^ varied in a pseudo-random fashion (50, 100, 200, and 500 ms)

^b^ varied in a pseudo-random fashion (25, 50, and 100 ms)

^c^ varied in a pseudo-random fashion (100, 200, 500, and 1000 ms)

^d^ varied in a pseudo-random fashion (250, 1000, and 2000 ms)

^e^ varied in a pseudo-random fashion (50, 100, and 200 ms)

* Driven high gamma (70-150 Hz) activity (13 samples)

^#^ Driven multiunit activity (3 samples)

Shaded in red: Pre-state power is significantly greater than Post-state power (Wilcoxon rank sum test, *P* < 0.05)

Shaded in blue: Post-state power is significantly greater than Pre-state power (Wilcoxon rank sum test, *P* < 0.05)

**Table S2.** Summary of the likelihood of detecting stimulation response to single microstimulation pulses between network nodes.

|  |  | Sender node |  |  |  |  |  |  |
| --- | --- | --- | --- | --- | --- | --- | --- | --- |
|  |  | OFC | CN | Put | dPFC | Amyg | BFB | NAc |
| Receiver node | OFC | 6/1358 | 4/615 | 0/94 | 0/51 | 6/59 | 0/6 | 0/25 |
|  | ACC | 7/1165 | 3/507 | 0/93 | 0/93 | 1/97 | 0/12 | 0/53 |
|  | CN | 14/1212 | 7/568 | 1/113 | 0/101 | 1/107 | 0/12 | 0/44 |
|  | Put | 1/180 | 5/112 | 0/26 | 1/25 | 1/29 | 1/3 | 0/14 |
|  | GPe | 1/92 | 1/49 | 0/9 | 0/11 | 1/12 | 0/2 | 0/3 |
|  | dPFC | 9/779 | 0/330 | 0/56 | 0/59 | 0/68 | 0/6 | 0/48 |
|  | M1 | 6/100 | 0/299 | 0/79 | 0/100 | 0/87 | 0/12 | 0/64 |
|  | S1 | 0/123 | 0/101 | 0/39 | 0/44 | 0/49 | 0/6 | 0/39 |
|  | PPC | 0/403 | 0/18 | 0/115 | 0/149 | 0/136 | 0/18 | 1/403 |
|  | Amyg | 0/29 | 0/21 | 0/8 | 0/9 | 0/6 | 0/1 | 0/4 |
|  | NAc | 0/11 | 0/9 | 0/1 | 0/6 | 0/2 |  |  |
|  | PrS | 0/15 | 0/12 | 0/3 | 0/5 | 0/4 | 0/1 | 0/1 |

Notes:

Notes:

<u>Cortical regions</u> ― OFC: orbitofrontal cortex; ACC: anterior cingulate cortex; dPFC: dorsal prefrontal cortex; M1: primary motor cortex; S1: primary somatosensory cortex; PPC: posterior parietal cortex

<u>Subcortical regions</u> ― CN: caudate nucleus; Put: putamen; GPe: globus pallidus external; Amyg: amygdala; BFB: basal forebrain; NAc: nucleus accumbens; PrS: presubiculum

Data are shown as N1/N2, where N1 is the number of single-pulse stimulation responding sites in each brain area and N2 is the total number of recorded sites in each brain area. Highlighted in red are statistically significant directed function interactions based on the binomial test.

**Table S3.** RSM motifs.

| Edge samples | Modulator |
| --- | --- |
| OFC→dPFC | ACC |
| OFC→CN | dPFC<br>ACC |
| OFC→CN | OFC |
| OFC→GPe | S1 |
| CN→CN | OFC |
| CN→CN | ACC |
| CN→CN | M1<br>PPC (SMG)<br>ACC<br>GPe<br>Put (2)<br>CN |
| CN→CN | PPC (pCun) (5)<br>S1 (2) |
| CN→OFC | GPe |
| CN→OFC | PCC<br>CN |
| CN→ACC | PCC (2) |
| CN→Put (SSM) | PPC (SPL)<br>PPC (SMG)<br>M1<br>CN |
| Amyg→OFC | PPC (SMG)<br>PPC (pCun) |
| Amyg→OFC | PPC (SMG)<br>CN |
| Amyg→OFC | PPC (SPL)<br>S1<br>ACC<br>PCC (2)<br>CN |
| Amyg→ACC | PPC (pCun)<br>Put |

**Table S4.**
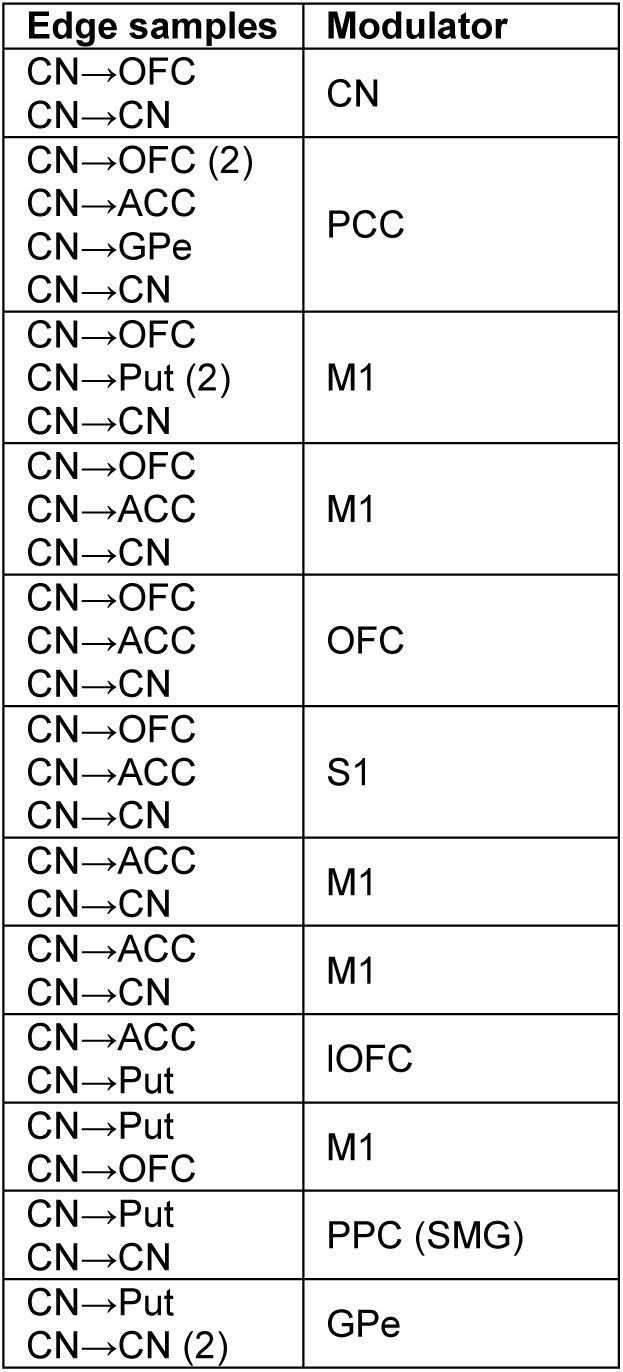
RNM motifs.

**Table S5.** SSM motifs.

| Edge samples | Modulator |
| --- | --- |
| CN→Put<br>BFB→Put | PPC (SPL)<br>PPC (SMG) (2)<br>PCC<br>M1 (3)<br>CN<br>GPe |

Notes:

Notes:

<u>Cortical regions</u> ― OFC: orbitofrontal cortex; ACC: anterior cingulate cortex; PCC: posterior cingulate cortex; dPFC: dorsal prefrontal cortex; M1: primary motor cortex; S1: primary somatosensory cortex; PPC: posterior parietal cortex (SPL: superior parietal lobule; SMG: supramarginal gyrus; pCun: precuneus)

<u>Subcortical regions</u> ― CN: caudate nucleus; Put: putamen; GPe: globus pallidus external; Amyg: amygdala; BFB: basal forebrain

